# Molecular taxonomy of human ocular outflow tissues defined by single cell transcriptomics

**DOI:** 10.1101/2020.02.10.942649

**Authors:** Gaurang Patel, Wen Fury, Hua Yang, Maria Gomez-Caraballo, Yu Bai, Tao Yang, Christina Adler, Yi Wei, Min Ni, Ying Hu, George Yancopoulos, W. Daniel Stamer, Carmelo Romano

**Author notes:** contributed equally to this work (Gaurang Patel, Wen Fury). Corresponding authors: Carmelo Romano, Ophthalmology Research, Regeneron Pharmaceuticals, Inc. 777 Old Saw Mill River Rd, Tarrytown, NY 10591-6707; W. Daniel Stamer, Departments of Ophthalmology and Biomedical Engineering, Duke University, DUMC 3802, Durham, NC, 27710.

## Abstract

The conventional outflow pathway is a complex tissue responsible for maintaining intraocular pressure (IOP) homeostasis. The coordinated effort of multiple cells with differing responsibilities ensure healthy outflow function and IOP maintenance. Dysfunction of one or more resident cell type results in ocular hypertension and risk for glaucoma, a leading cause of blindness. In this study, single cell RNA sequencing was performed to generate a comprehensive cell atlas of human conventional outflow tissues. We obtained 17757 genes expression profiles from 8758 cells from eight eyes of four donors representing the outflow cell transcriptome. Upon clustering analysis, 12 distinct cell types were identified, and region-specific expression of candidate genes were mapped in human tissues. Significantly, we identified two distinct expression patterns (myofibroblast and fibroblast) from cells located in the trabecular meshwork (TM), the primary structural component of the conventional outflow pathway. We also located neuron and macrophage signatures in the TM. The second primary component structure, Schlemm’s canal displayed a unique combination of lymphatic/blood vascular gene expression. Other expression clusters corresponded to cells from neighboring tissues, predominantly in the ciliary muscle/scleral spur, which together correspond to the uveoscleral outflow path. Importantly, the utility of our atlas was demonstrated by mapping glaucoma-relevant genes to outflow cell clusters. Our study provides a comprehensive molecular and cellular classification of conventional and unconventional outflow pathway structures responsible for IOP homeostasis.

**Significance statement:** Ocular hypertension is the primary, and only modifiable risk factor for glaucoma, the leading cause of irreversible blindness. Intraocular pressure is regulated homeostatically by resistance to aqueous humor outflow through an architecturally complex tissue, the conventional/trabecular pathway. In this study, we generated a comprehensive cell atlas of the human trabecular meshwork and neighboring tissues using single cell, RNA sequencing. We identified 12 distinct cell types, and mapped region-specific expression of candidate genes. The utility of our atlas was demonstrated by mapping glaucoma-relevant genes to conventional outflow cell clusters. Our study provides a comprehensive molecular and cellular classification of tissue structures responsible for intraocular pressure homeostasis in health, and dysregulation in disease.

## INTRODUCTION

Elevated intraocular pressure (IOP) is a major causative risk factor for the development(1) and progression(2, 3) of most forms of glaucoma. Efficacious lowering of IOP, whether elevated or not slows glaucomatous disease progression(2, 4). Current medical treatments lower IOP by three different mechanisms: suppressing aqueous humor formation, or increasing its rate of outflow via the unconventional or conventional outflow pathways (5, 6). While all three target tissues participate in aqueous humor dynamics, the conventional (trabecular) outflow pathway is responsible for homeostatically regulating IOP by the coordinated generation of outflow resistance involving cells that reside in the trabecular meshwork (TM) and Schlemm’s canal (SC)(7, 8) (9, 10). Importantly, dysfunction in the regulation of conventional outflow resistance in results in elevated IOP in glaucoma patients(11, 12).

The TM is an avascular, complex connective tissue located at the iridocorneal angle, bridging from Schwalbe’s line anteriorly to the scleral spur/ciliary muscle posteriorly. Anatomically the TM is divided into three distinct tissue layers, the inner uveal meshwork, the middle corneoscleral meshwork, and the outer juxtacanalicular tissue (JCT, also known as cribriform tissue). Functioning as a biological filter, the uveoscleral and corneoscleral meshwork consists of collagen and elastin lamellae/plates covered by TM cells. In contrast, the JCT is comprised of TM cells embedded in the extracellular matrix of a loose connective tissue, directly interfacing with the inner wall of SC (reviewed by Stamer and Clark, 2016; Tamm, 2009)(7, 8). Cells that populate the TM are all of neural crest in origin(13), displaying different morphologies depending upon their tissue location. Thus, TM cells in uveal/ corneoscleral meshwork display endothelial and macrophage properties, maintaining patent flow passageways by secreting antithrombotic molecules(14), phagocytosing cellular debris, neutralizing reactive oxygen species(15–17), and mediating immune function(18, 19). While the cells in JCT region display fibroblast and smooth-muscle like properties, playing an important role in the generation and control of outflow resistance(7). In response to various stimuli including mechanical cues(20–22), JCT cells continuously repair/ remodel the extracellular matrix(23, 24) and maintain contractile tissue tone in conjunction with the ciliary muscle. The importance of TM contractile tone in IOP regulation has been recently exploited pharmacologically by a new class of glaucoma drugs, the rho-kinase inhibitors which selectively relax the TM, and decrease outflow resistance (25).

Studies involving the use of TM and SC cells in cell and organ culture has resulted the identification of multiple drug targets that are in various stages of clinical development, including rho kinase inhibitors(26). However, due to their multiple responsibilities and unique environment, specific markers have not been identified for conventional outflow cells. Instead, cell morphology, growth characteristics, behavior, dissection technique and a panel of protein markers are used for their identification and characterization. Typical panel of markers used to identify TM cells includes chitinase-3 like-1, matrix GLA protein (MGP), aquaporin-1(26–29), and glucocorticoid induced upregulation of myocilin (MYOC) (30). Several transcriptomic and proteomic studies have been conducted to identify marker proteins, however they were all based upon bulk tissue preparations that failed to identify cell-specific markers (31–36).

Recent advances in single-cell RNA sequencing allows molecular separation of different cell populations based on expression profile at the single cell level(37, 38). Such studies have been conducted using ocular tissues of the whole retina (39–41), limbal/ corneal epithelia tissues(42), but not outflow tissues. Like the retina, outflow tissues are ideal tissue to study at single cell level because of their complex architecture and variety of cell types. Significantly, the critical role that TM and SC cells play in outflow resistance/IOP regulation and their accessibility for therapeutic interventions makes them attractive targets for gene therapy for both congenital and adult forms of glaucoma. Moreover, identification of TM/SC cell-specific expression of proteins is critical for identification of tissue-specific promoters needed for mechanistic studies involving transgenic/knockout mice.

To address these current deficiencies holding back glaucoma research, we performed single cell RNA-seq of outflow tissues from human donor eyes and have identified cells in human outflow tissues that have 12 unique molecular signatures. Locations of cells having these RNA profiles were mapped using in situ hybridization and immunohistochemistry of human eye tissue. Taken together, this study provides a comprehensive molecular and cellular classification of conventional and unconventional outflow pathway tissues responsible for IOP regulation.

## RESULTS

### Single cell transcriptome atlas of human outflow cell types

To generate an atlas of human conventional outflow tissues, we isolated TM tissue from 8 different human donor eyes using a blunt dissection (Figure S1A, S1B), and subjected the dissected tissue to enzymatic digestion to prepare single cell suspension. Single cell suspensions were checked for cell viability, with cells from 4 donor pairs (8 samples) included in subsequent droplet-based single cell RNA sequencing (sc-RNAseq) using 10x genomics platform. Using Seurat 2.3 software, the average number of total UMIs detected were 5000 (Figure S2A, S3A), average gene counts were 2000 (Figure S2B, S3B) and the average mitochondrial gene count was less than 0.2% (figure S2C, S3C). After the QC metrics described in the method section were applied, 17,757 genes and 8,758 cells were used in clustering analyses. Each of the human TM samples contributed uniformly to all clusters (Figure S4) and the percentage of cells in each cluster is registered in Table 2, except for lymphatic-like endothelial cells and epithelium cells, which had low percentage in the overall cell population (1.42% and 0.34%, respectively). Twelve cell clusters were identified using the Seurat algorithm. Their identifies were discovered by using cluster specific genes as well as canonical cell type markers (Figure 1A and 1B). From the most abundant to the least, cell signatures included Schwann cell like, TM1-fibroblast like, smooth muscle cell, TM2-myofibroblast like, melanocyte, macrophages, pericyte, vascular endothelium cell, T/NK cell, lymphatic-like endothelium, myelinating Schwann cell and epithelium cell clusters (Table 2). To better characterize each cell type, we studied the similarity and dissimilarity of the cell types by using average UMI of each gene across all cells within the same cell type for hierarchical clustering analysis. As expected, our results indicate that TM1-fibroblast-like and TM2-myofibroblast-like cells are similar to each other. Likewise, macrophage were similar to T/NK cells, pericytes shared likeness with smooth muscle and vascular resembled lymphatic endothelial cells. Lastly, melanocytes, which could derive from Schwann cells during development, are similar to Schwann cells or Schwann cell-like cells (Figure 1C).

**Figure 1:**
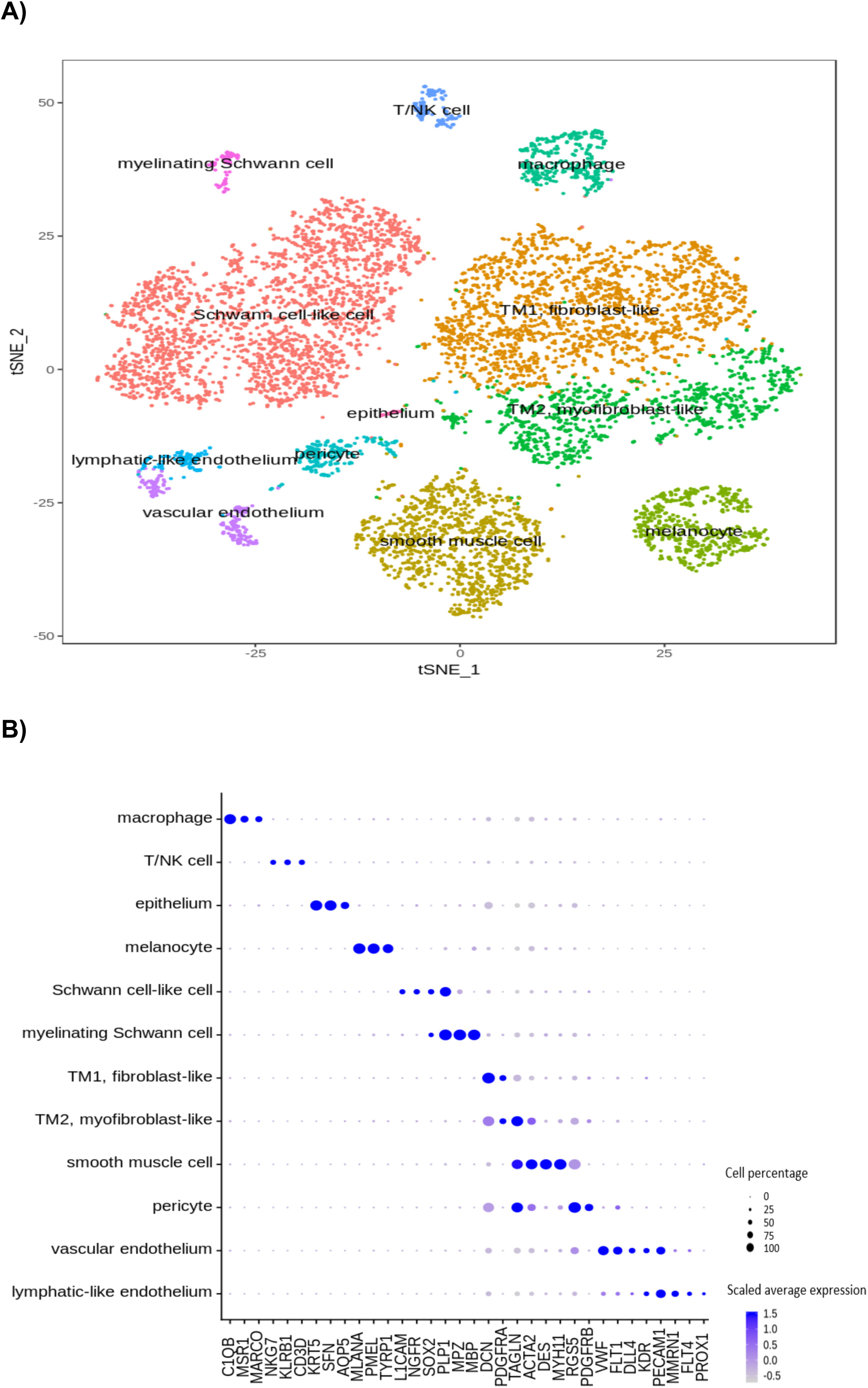
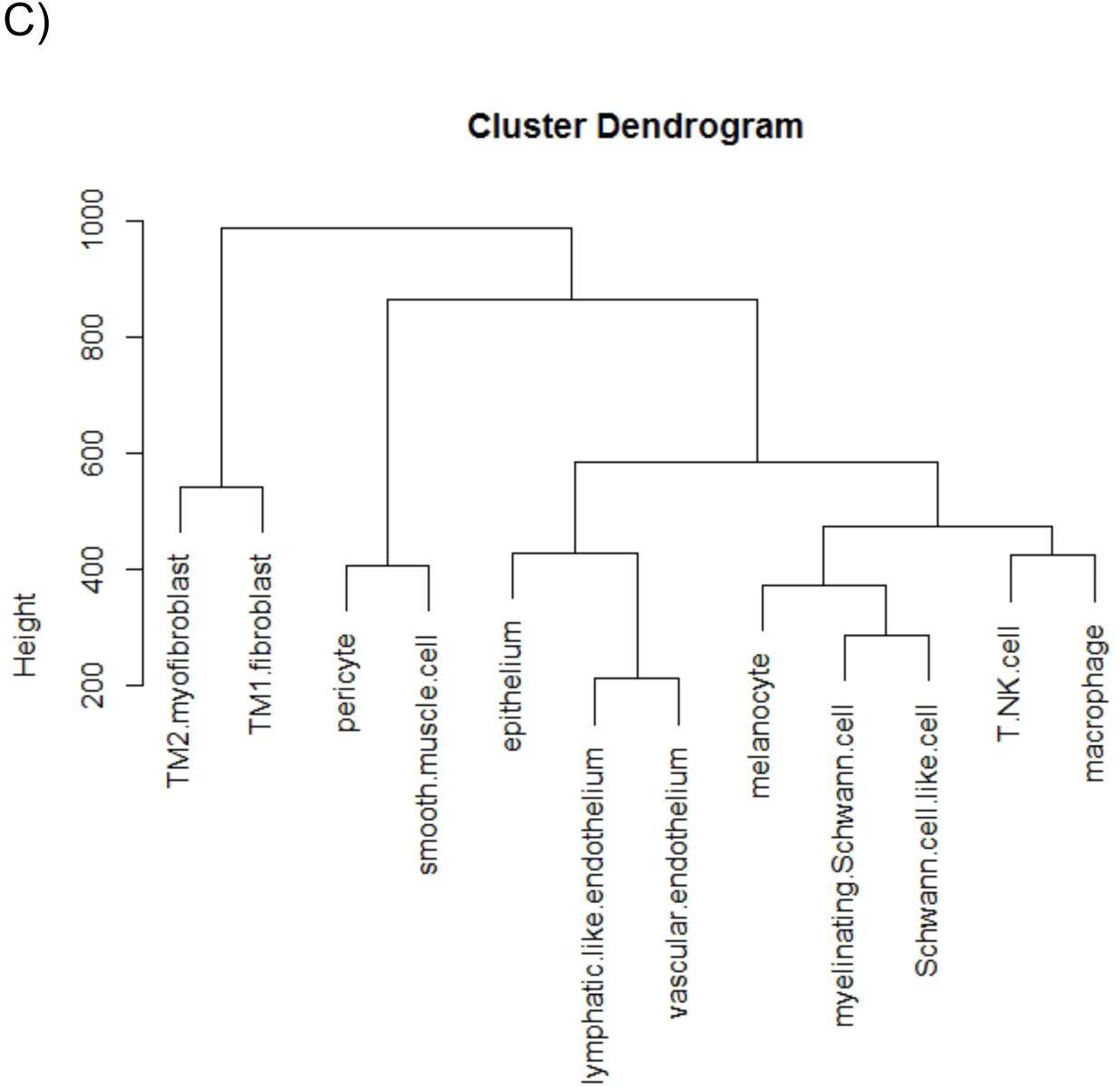
Identification of human outflow cell types. **A)** T-distributed Stochastic Neighbor Embedding (tSNE) visualization of outflow tissue transcriptome heterogeneity of 8758 cells. 12 distinct cell clusters were identified in the outflow tract, and cells are colored and labelled by cluster assignments. **B)** Dot plot of gene expression combinations that uniquely identified each of the cell clusters. Quality control metrics are described in methods section. The size of each circle is proportional to the percentage of cells expressing the gene (marker) within the cluster and its intensity depicts the average transcript count within expressing cells. **C)** Hierarchical clustering of cell types, showing similarity or dissimilarity among cell types.

**Table 2:** Cell numbers and percentage by cell types and sample contribution.

|  | hTM_101 | hTM_102 | hTM_103 | hTM_104 | hTM_107 | hTM_108 | hTM_109 | hTM_110 | sum | percent |
| --- | --- | --- | --- | --- | --- | --- | --- | --- | --- | --- |
| Schwann cell-like cell | 394 | 334 | 270 | 208 | 241 | 519 | 315 | 242 | 2523 | 28.81 |
| TM1, fibroblast-like | 189 | 207 | 242 | 138 | 335 | 570 | 278 | 192 | 2151 | 24.56 |
| smooth muscle cell | 44 | 105 | 385 | 68 | 76 | 356 | 64 | 90 | 1188 | 13.56 |
| TM2, myofibroblast-like | 95 | 89 | 83 | 92 | 277 | 70 | 147 | 242 | 1095 | 12.50 |
| melanocyte | 9 | 3 | 15 | 7 | 128 | 246 | 118 | 66 | 592 | 6.76 |
| macrophage | 9 | 22 | 24 | 42 | 28 | 31 | 102 | 103 | 361 | 4.12 |
| pericyte | 29 | 29 | 13 | 25 | 48 | 42 | 28 | 24 | 238 | 2.72 |
| vascular endothelium | 66 | 41 | 10 | 15 | 27 | 26 | 18 | 5 | 208 | 2.37 |
| T/NK cell | 33 | 22 | 9 | 23 | 14 | 14 | 18 | 13 | 146 | 1.67 |
| lymphatic-like endothelium | 33 | 44 | 10 | 19 | 17 | 0 | 0 | 1 | 124 | 1.42 |
| myelinating Schwann cell | 8 | 3 | 10 | 5 | 19 | 9 | 27 | 21 | 102 | 1.16 |
| epithelium | 2 | 0 | 15 | 6 | 6 | 1 | 0 | 0 | 30 | 0.34 |
| sum | 911 | 899 | 1086 | 648 | 1216 | 1884 | 1115 | 999 | 8758 | 100.00 |

### Characterization and localization of cluster specific cell markers in human TM by RNAScope

We found genes that were enriched and sometimes specific to each cluster beyond canonical cell-type-specific markers. To locate each of the clusters identified by scRNAseq in human outflow tissues we performed RNAScope with different gene-specific probes. For identifying Schwann cell like cluster, we selected RNA probes selective for Aquaporin 7 Pseudogene 1 (AQP7P1) (Figure 2A) and Sodium Voltage-Gated Channel Alpha Subunit 7 (SCN7A) (Figure 2B). These markers localized to scleral spur and ciliary muscle regions. We observed that Decorin (DCN) (Figure 3A) and platelet-derived growth factor receptor alpha (PDGFRA) (Figure 3B) were highly enriched in TM1-fibroblast cell cluster, but interestingly localized to several tissues including scleral spur, the JCT and SC regions of TM. In contrast, R-spondin-4 (RSPO4) (Figure 4A) which were more specific to TM2-myofibroblast like cluster, mapped preferentially to throughout the TM. Whereas its homologue, R-spondin-2 (RSPO2) (Figure 4B) was abundant in both TM clusters, being expressed throughout the TM and neighboring tissues.

**Figure 2:**
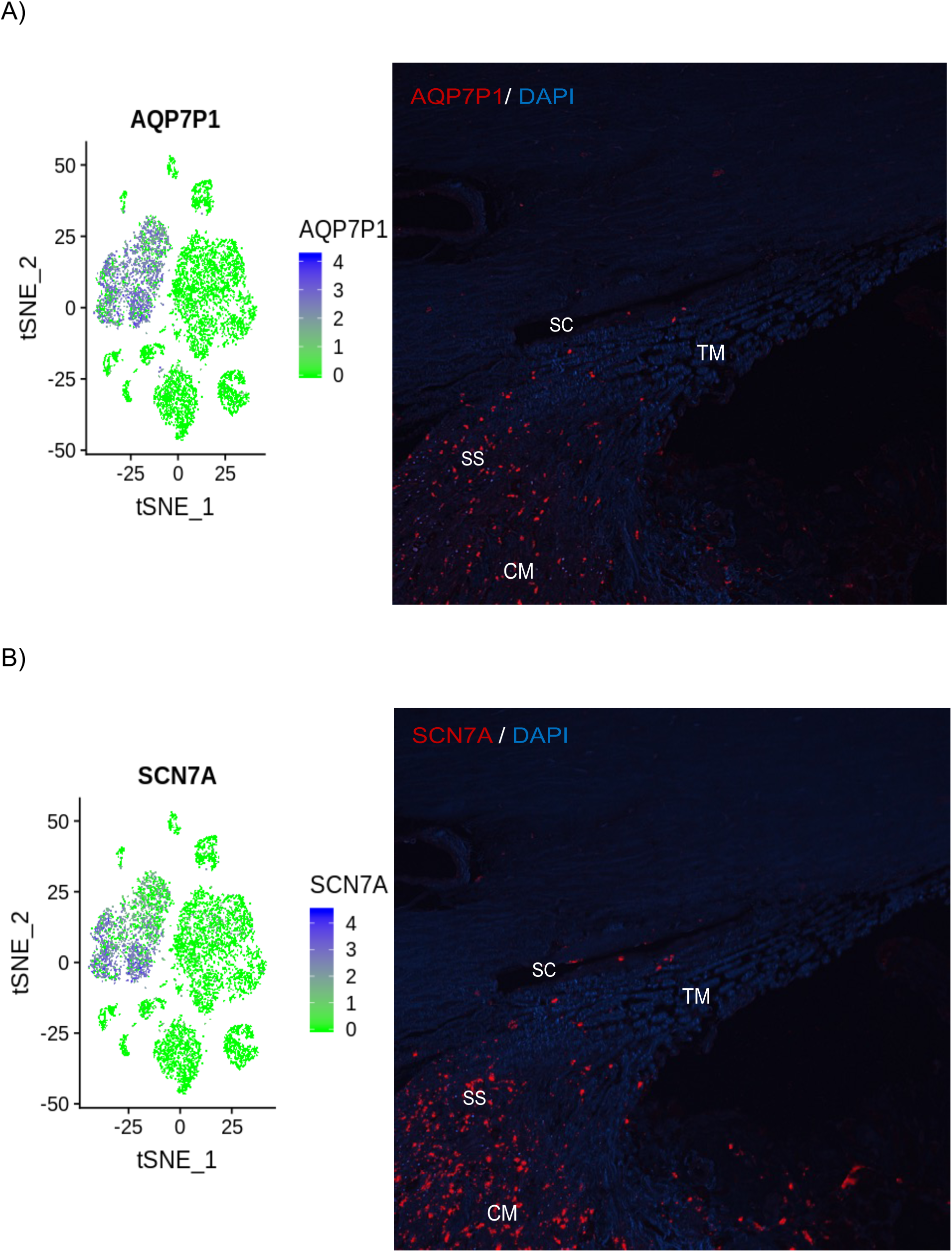
In situ hybridization mapping of cells from schwann cell like cluster in human eye sections. A t-distributed stochastic neighbor embedding (tSNE) plot showing relative expression of gene (blue dots) in each cluster is displayed on the left, and on the right is ISH stained human eye section showing mRNA signal (red fluorescence) from gene of interest. mRNA probes corresponding to cluster markers, AQP7P1 **(A)** and SCN7A **(B)** predominantly localized to the scleral spur and ciliary muscle regions, with some cells extending into the TM. DAPI staining (blue) counterstains cell nuclei. Magnification: 20x. TM- Trabecular meshwork; CM- ciliary muscle; SC- Schlemm’s canal; SS- Scleral spur. Scale in tSNE plot shows intensity of the average transcript count of particular marker within expressing cells and clusters.

**Figure 3:**
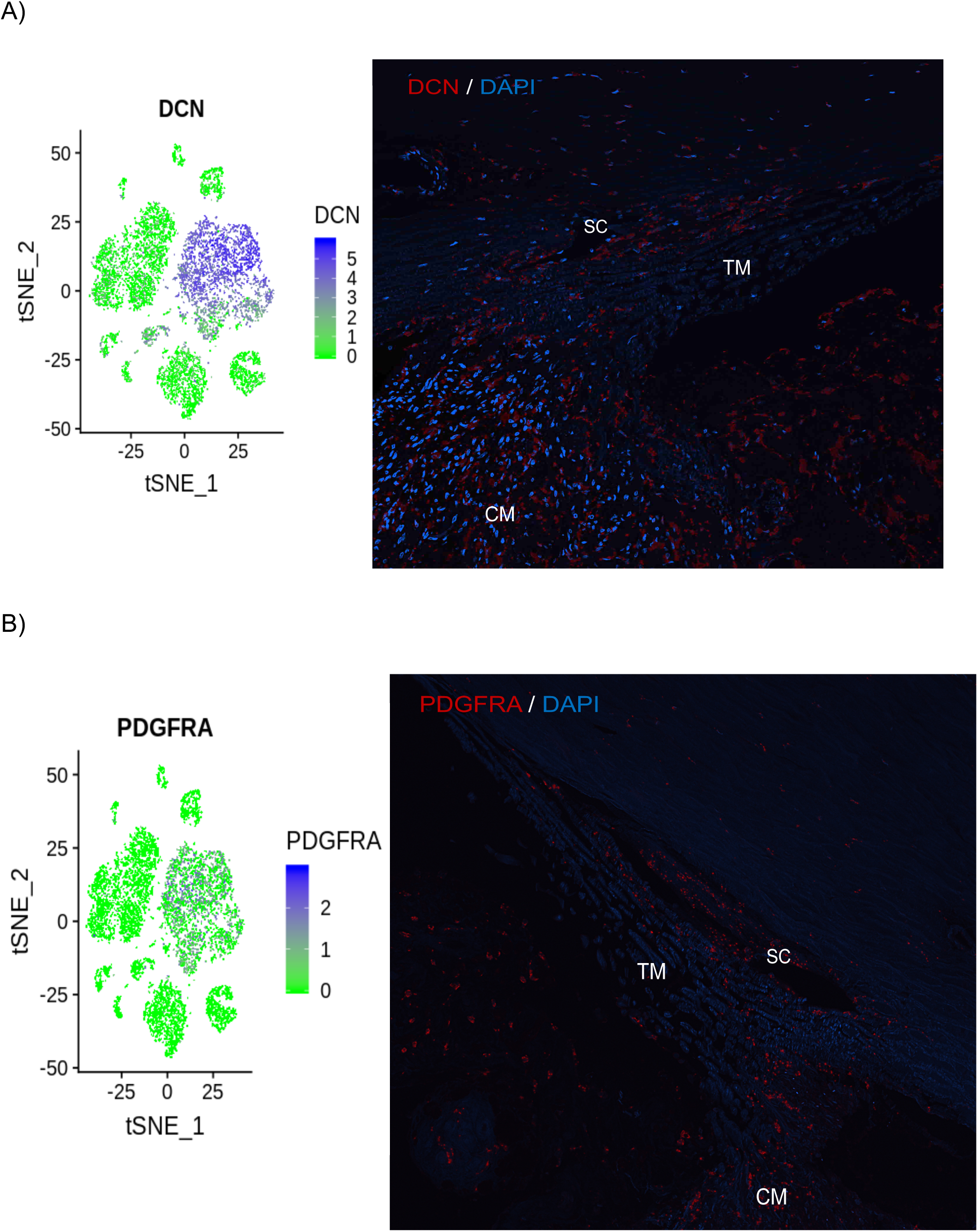
Localization of the TM1- fibroblast like cell clusters using in situ hybridization of human eye sections. The left side shows a t-distributed stochastic neighbor embedding (tSNE) plot of relative gene of interest expression (blue dots) in each cluster, while the right-side displays ISH stained human eye section mapping mRNA signal as red fluorescence. mRNA probes specific for cluster markers, DCN **(A)** and PDGFRA **(B)** were localized predominantly to juxtacanalicular tissue (JCT) and SC regions of conventional tract, however scleral spur cells also were also labeled. DAPI staining (blue) counterstains cell nuclei. Magnification: 20x. TM- Trabecular meshwork; CM- ciliary muscle; SC- Schlemm’s canal. Scale in tSNE plot shows intensity of the average transcript count of particular marker within expressing cells and clusters.

**Figure 4:**
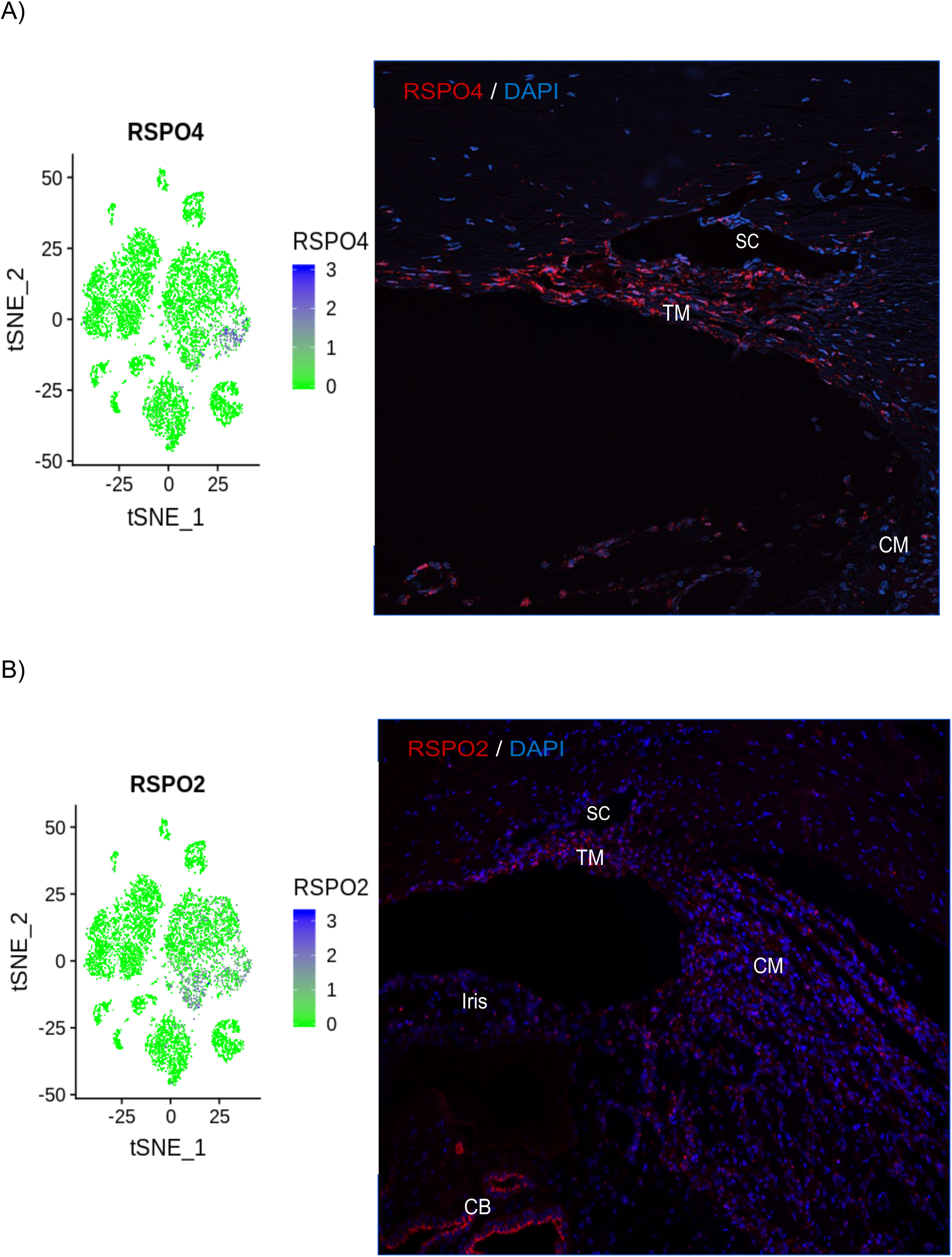
Mapping of gene candidates from TM2- myofibroblast like cell cluster using in situ hybridization of human eye sections. For each image, the left side is a t-distributed stochastic neighbor embedding (tSNE) plot showing relative expression of gene of interest (blue dots) in each cluster and right-side is ISH stained human eye section showing mRNA signal as red fluorescence. mRNA probes to candidates, RSPO4 **(A)** and RSPO2 **(B)** were tested. RSPO4 expression was spread throughout the TM, but not surrounding tissues. Whereas, RSPO2 expression was localized throughout the TM and neighboring tissues. DAPI staining (blue) counterstains cell nuclei. Magnification: 20x for figure 4A and 10x for figure 4B. TM- Trabecular meshwork; CM- ciliary muscle; SC- Schlemm’s canal; CB- ciliary body. Scale in tSNE plot shows intensity of the average transcript count of particular marker within expressing cells and clusters.

We found the smooth muscle cluster gene muscle myosin heavy chain 1 (MYH11) (Figure 5A) limited to ciliary muscle region. Transgelin (TAGLN) (Figure 5B) was highly enriched in TM2 and smooth muscle cluster, correspondingly localized to both TM and ciliary muscle. Lymphatic Vessel Endothelial Hyaluronan Receptor 1 (LYVE1) (Figure 6A), C1QB (Figure 6B), TYRO Protein Tyrosine Kinase Binding Protein (TYROBP) (Figure 6C) were highly enriched in the macrophage cluster, and RNAScope revealed focal, macrophage-like labeling spread throughout the conventional outflow pathway. These results show that only three of the bioinformatically identified cell types (TM1, TM2, macrophages) are found in the anatomically defined TM, and the others reflect cells from neighboring tissues (i.e.: the unconventional outflow pathway).

**Figure 5:**
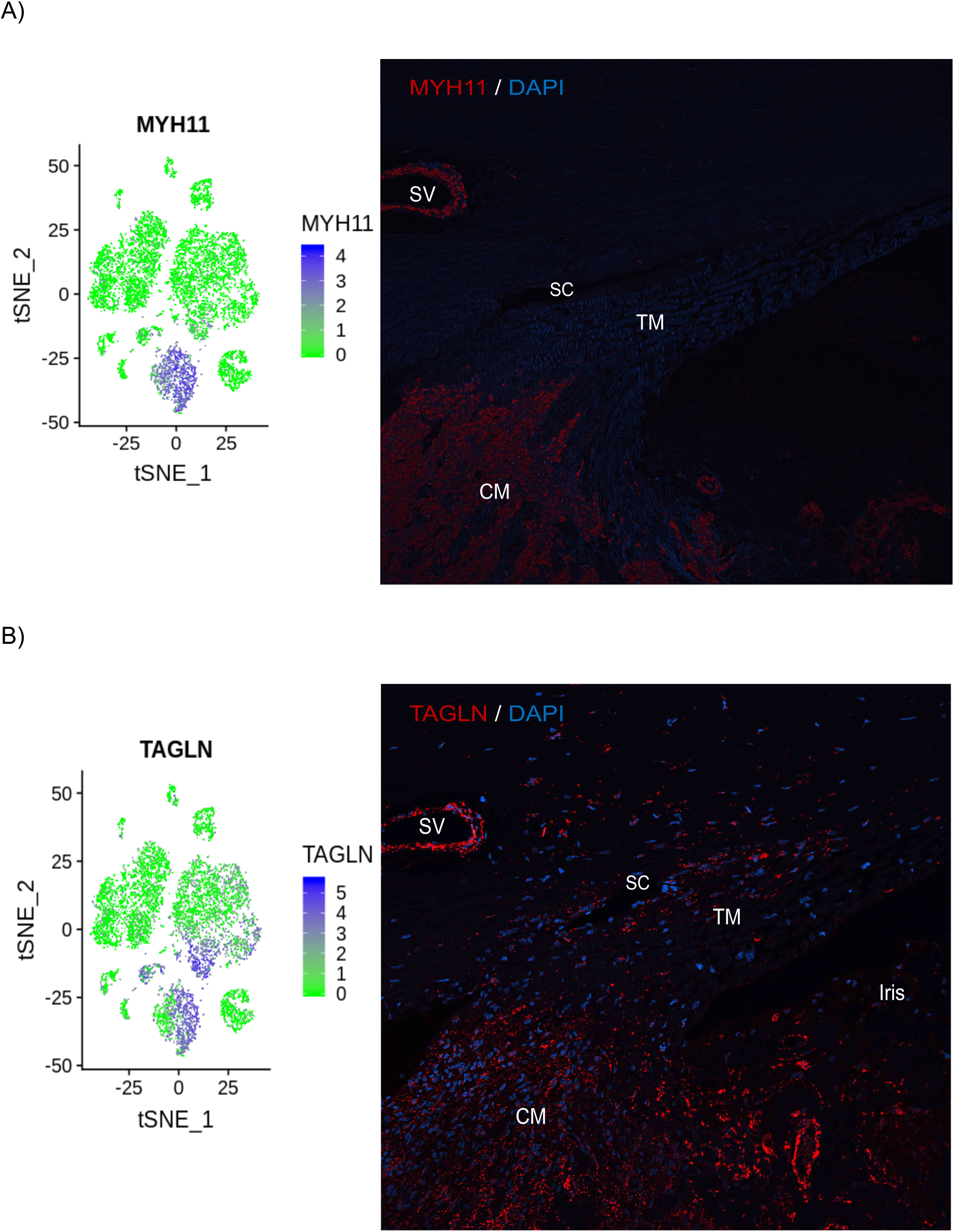
Localization of smooth muscle cell cluster gene candidates using in situ hybridization of human eye sections. On the left side are t-distributed stochastic neighbor embedding (tSNE) plots showing relative candidate gene expression (blue dots) in each cluster and on the right-side are ISH stained human eye sections showing specific mRNA signal as red dot fluorescence. mRNA probes to MYH11 **(A)** and TAGLN **(B)** were predominantly confined to ciliary muscle, however TAGLN expression extended to a few cells in TM that were proximal to ciliary muscle. DAPI staining (blue) counterstains cell nuclei. Magnification: 20x. TM- Trabecular meshwork; CM- ciliary muscle; SC- Schlemm’s canal; SV- Scleral vessel. Scale in tSNE plot shows intensity of the average transcript count of particular marker within expressing cells and clusters.

**Figure 6:**
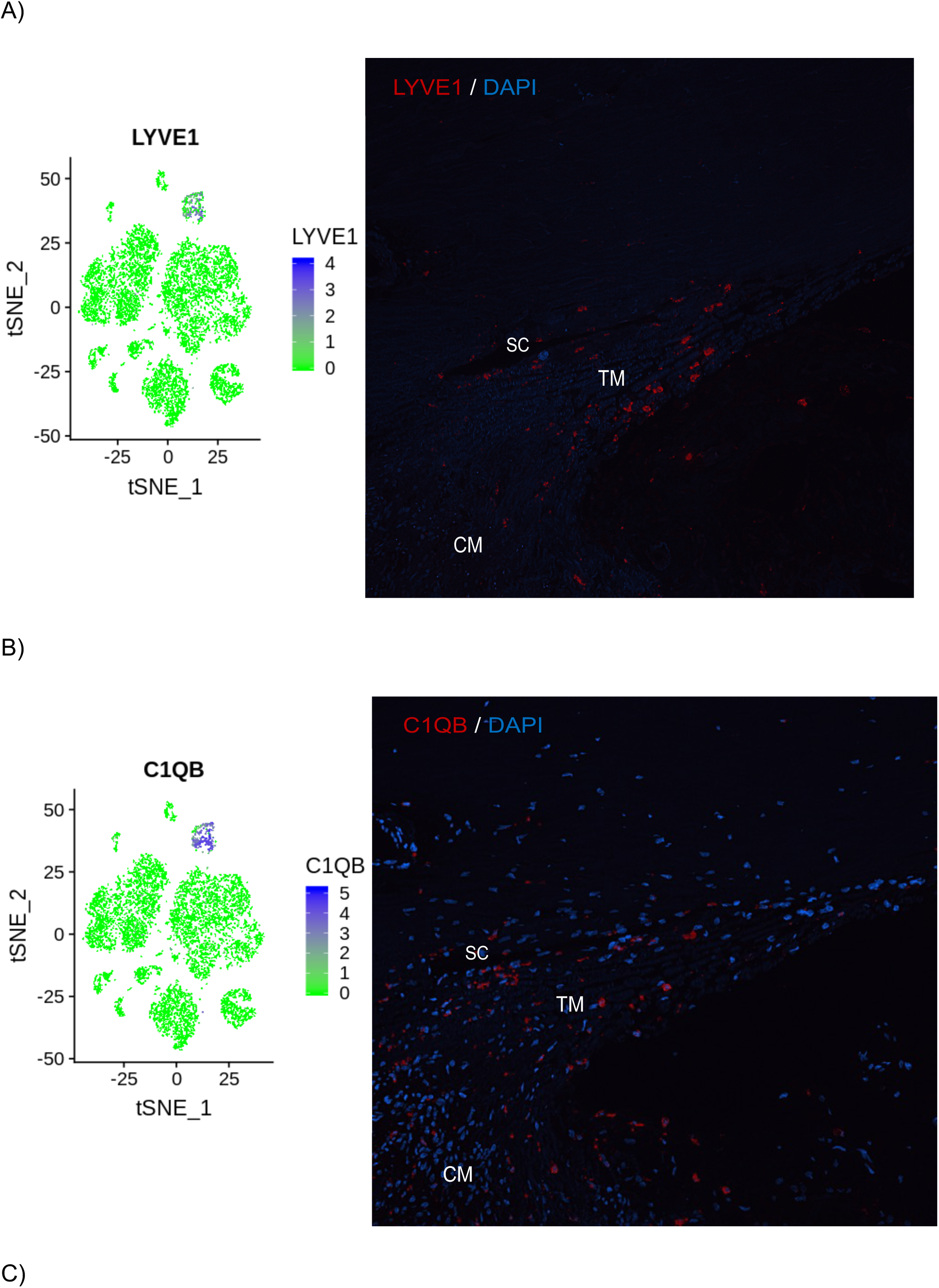
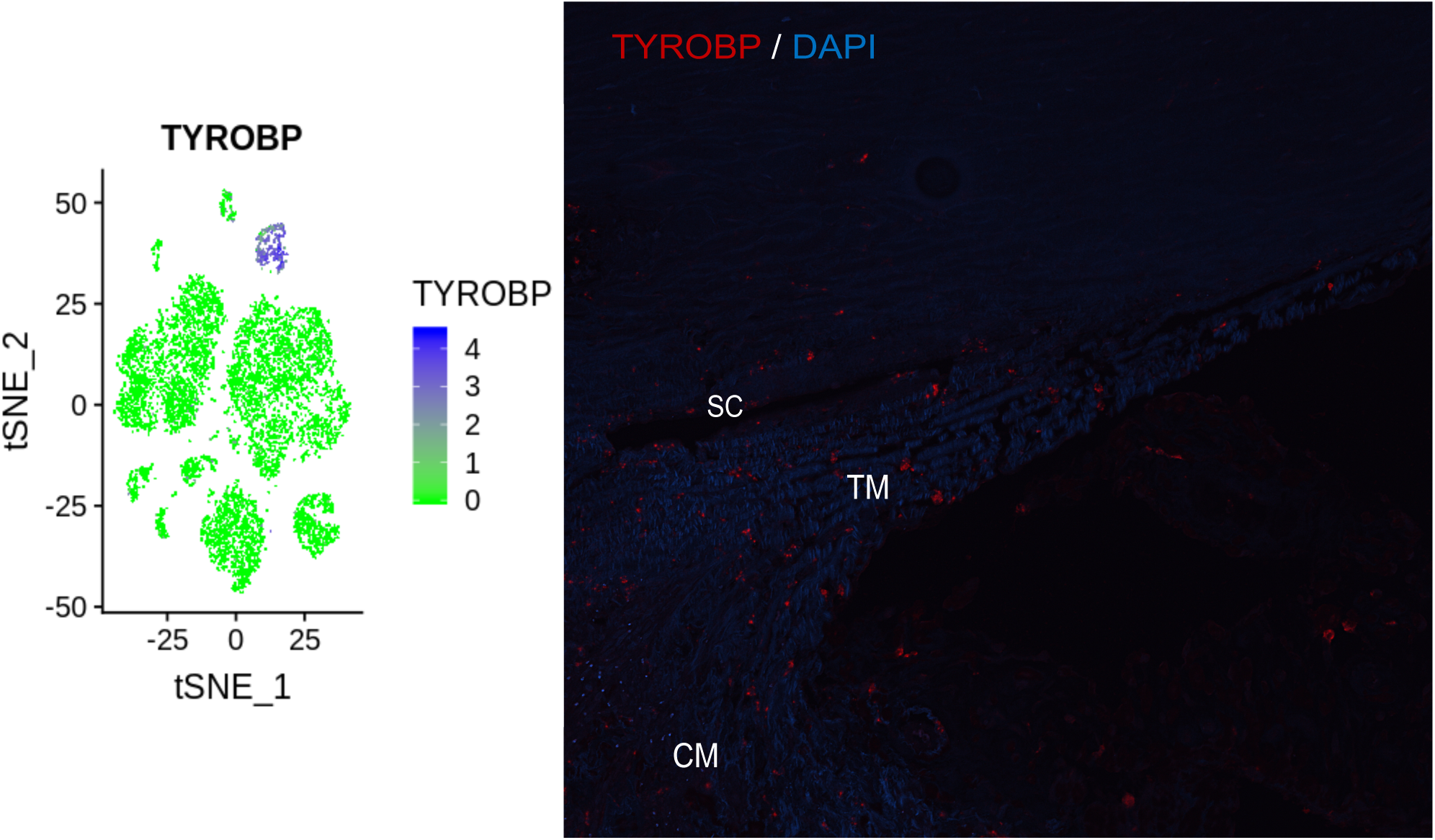
Mapping of macrophage genes in human outflow tissues using in situ hybridization. For each panel, the left side is t-distributed stochastic neighbor embedding (tSNE) plot showing relative expression of gene of interest (blue dots) in each cluster and right-side is ISH stained human eye section showing mRNA signal as red dot fluorescence. mRNA probes to LYVE1 **(A)**, C1QB **(B),** and **(C)** TYROBP show that macrophages are present throughout the TM, ciliary muscle and around SC. DAPI staining (blue) counterstains cell nuclei. Magnification: 20x. TM- Trabecular meshwork; CM- ciliary muscle; SC- Schlemm’s canal. Scale in tSNE plot shows intensity of the average transcript count of particular marker within expressing cells and clusters.

### Lymphatic-like endothelial cell cluster have both lymphatic and vascular endothelial phenotypes

Schlemm’s canal (SC) plays an important role in IOP regulation via synergistic interaction with JCT cells (25, 43). SC is unique structure displaying both blood vascular and lymphatic characteristics(44). Our scRNAseq data confirms these findings, with our transcriptome atlas showing a lymphatic-like endothelial cluster. Upon in-depth analysis we found that this cluster expressed markers for both vascular and lymphatic endothelial like cells (Figure 7). FLT4, FLT1, and Fibronectin (FN1) (Figure 8A, 8B, 8C) were expressed in the lymphatic-vascular endothelium cells and they were all found to be in the SC region. Although FN1, FLT1 was highly expressed in lymphatic endothelial and vascular endothelial clusters respectively, they were also present in other clusters. FN1 was localized more to SC and JCT regions and enriched in scleral blood vessels. FLT1 was localized more in SC and ciliary muscle. In addition, our single cell transcription analysis also provided the expression profiles of these unique cells at the whole genome transcriptome level.

**Figure 7:**
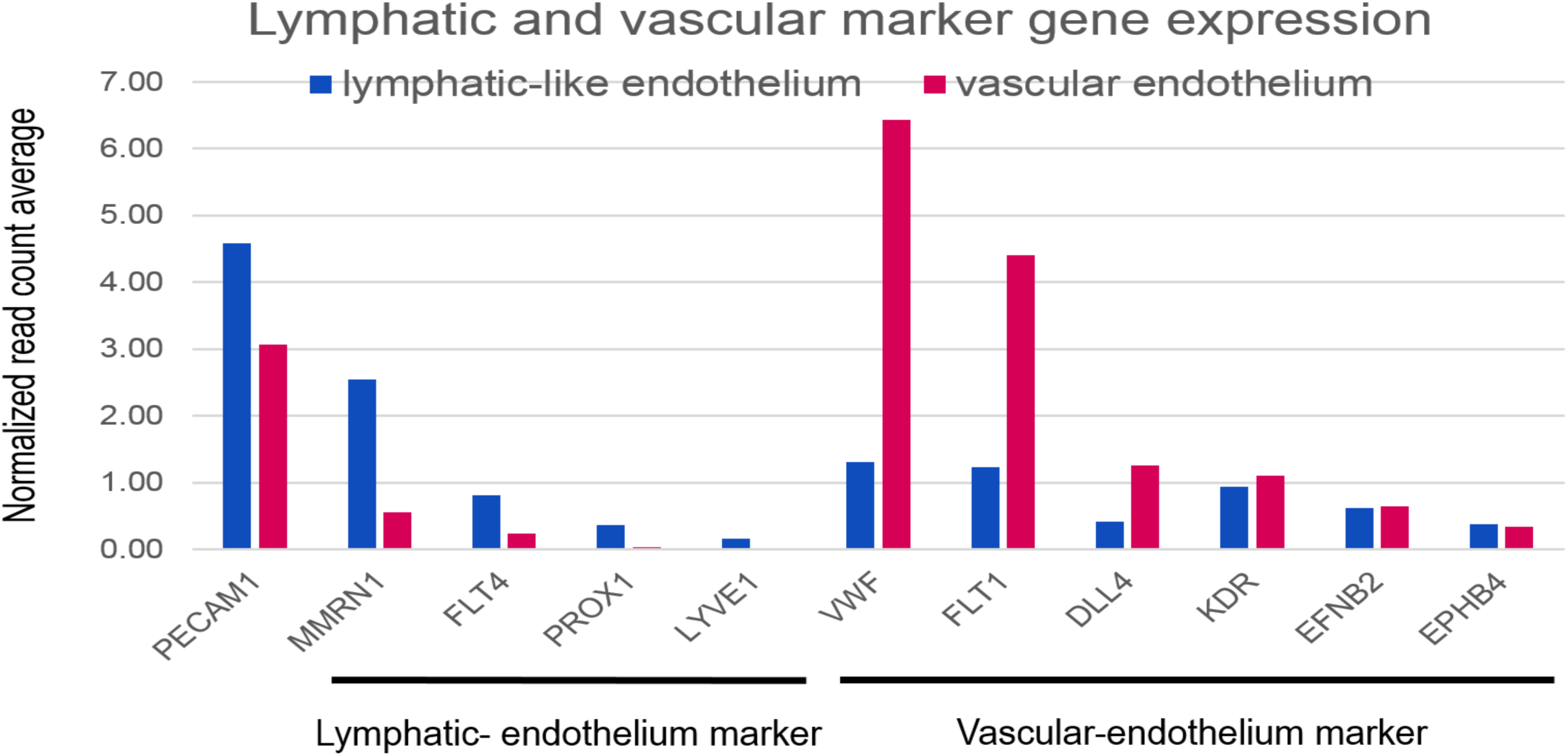
Lymphatic and vascular marker expression in scRNAseq outflow tissue transcriptome. Lymphatic-like endothelial cells express the pan endothelial marker PECAM1 and also express higher levels of lymphatic endothelial cell markers, MMRN1 and FLT4 than vascular endothelial cells, but lower levels of vascular endothelial cell markers, VWF and FLT1.

**Figure 8:**
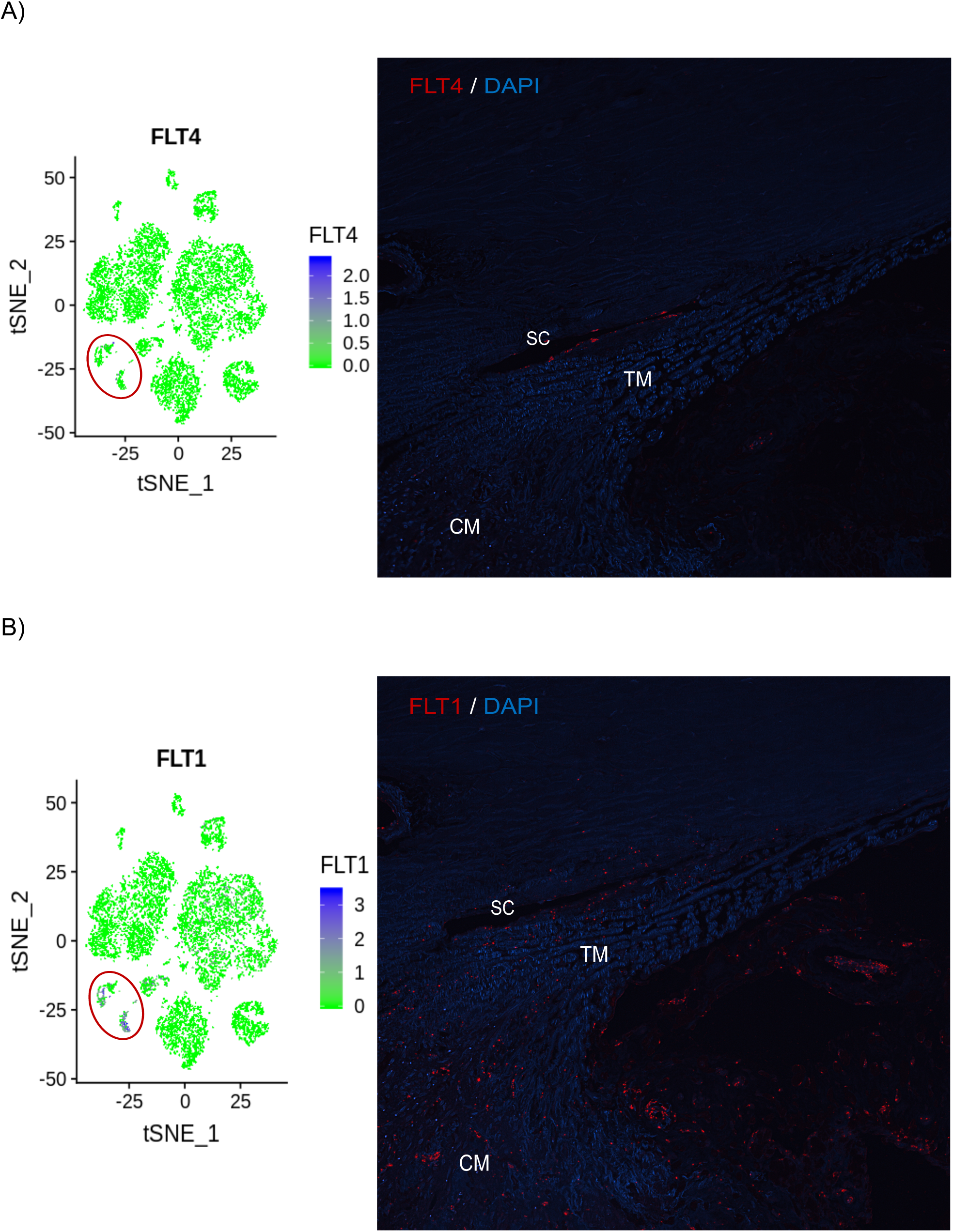
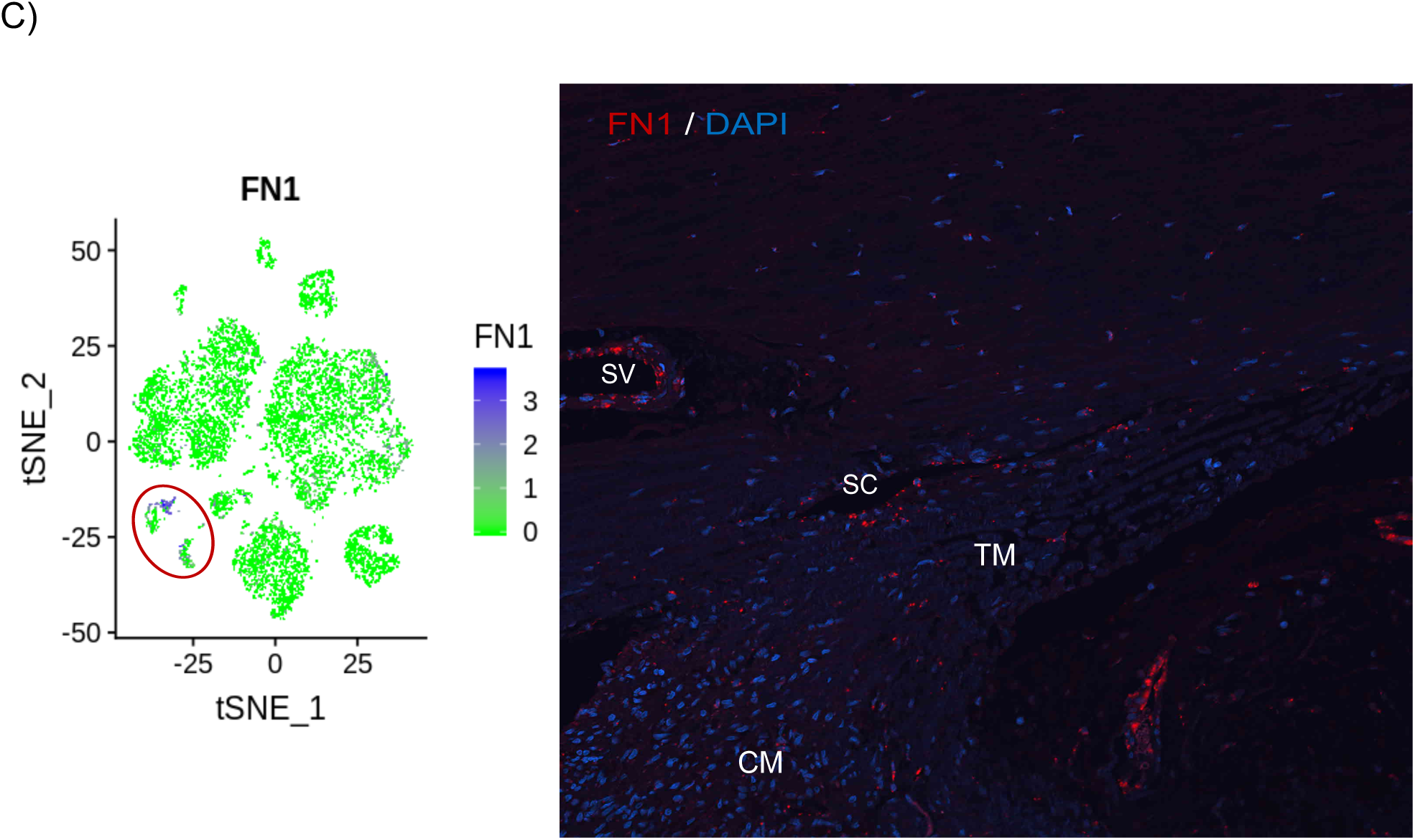
Localization of lymphatic and vascular endothelial cell expression in human eye sections using in situ hybridization. For each panel, the left side is a t-distributed stochastic neighbor embedding (tSNE) plot showing relative expression of candidate gene (blue dots) in each cluster and the right-side is stained human eye section showing mRNA signal as red fluorescence. mRNA probes corresponding to FLT4 **(A)**, FLT1 **(B),** and **(C)** FN1 were predominantly found in the SC region. FN1 was localized more to SC and JCT regions and FLT1 was localized more in SC, ciliary muscle and scleral vessel in human eye sections. DAPI staining (blue) counterstains cell nuclei. Magnification: 20x. TM- Trabecular meshwork; CM- ciliary muscle; SC- Schlemm’s canal. SV: scleral vessel. White circle represents Schlemm’s canal. Scale in tSNE plot shows intensity of the average transcript count of particular marker within expressing cells and clusters. Red circle in tSNE plot highlights lymphatic-vascular endothelial cell clusters.

### Cell type specific expression of glaucoma-related genes

There are many genes that are implicated or associated with elevated IOP and glaucoma. We selected few to see in which cell cluster they are present (Table 3; Figure 9A, 9B, 9C). Mutations in Myocilin (MYOC) cause glaucoma and Angiopoietin like 7 (ANGPTL7) (Figure 9A, 9B) is elevated in aqueous humor of glaucoma patients (45–47). MYOC was highly expressed TM1, TM2, and smooth muscle clusters and it was expressed at high levels in TM and at lower levels in the ciliary muscle, SC and scleral fibroblasts. Similarly, ANGPTL7 was found in both TM1 and TM2 clusters, but its expression was more limited, localizing to JCT and SC. Polymorphisms in a locus containing two genes, Caveolins CAV1 and CAV2 are also implicated in elevated IOP and glaucoma(48, 49). In our scRNAseq outflow transcriptome, CAV1 and CAV2 were present in various different clusters including lymphatic-vascular endothelial, pericyte, smooth muscle cell, myelinating schwann cell, melanocyte, epithelium cell clusters (Figure 9C). Interestingly, CAV1 and CAV2 were expressed at low levels in TM1 and TM2, but highly expressed in lymphatic-vascular endothelia cell cluster. Autotaxin (gene name: ENPP2) has been shown to be elevated in aqueous humor of glaucoma patients(50, 51) and studies have shown that autotaxin inhibitors lower IOP in mice and rabbits(50, 52). In the outflow transcriptome, ENPP2 was expressed in schwann cell like and melanocyte cell clusters (Figure 9C). Polymorphisms in the NOS3 gene encoding the endothelial-specific isoform of NOS (eNOS) impart risk for ocular hypertension and glaucoma(53–56). Data here show that indeed, NOS3 expression in the conventional outflow tract is confined to vascular endothelial/lymphatic cell clusters. Both Tie2 and Angpt1 loss of function variants associate with risk of congenital glaucoma, and single-nucleotide polymorphisms in the Angpt1 promoter region significantly associate with primary open-angle glaucoma risk(57–59). Consistent with immunofluorescence data in mice(60, 61), our ssRNAseq data show that expression of Angpt1 in the conventional outflow tract is limited to TM cell clusters, while Tek (Tie2) and its antagonist, Angpt2 are expressed by lymphatic/endothelia cell clusters. Many studies have described use of chitinase-3 like-1, matrix GLA protein (MGP), aquaporin-1, *α*B-crystallin (CRYAB) (7, 26–29, 62) as markers to identify TM cells. In our scRNAseq transcriptome, CHI3L1 was confined to TM2 and TM1 cell clusters, MGP to localized to all cell clusters with more preference to TM1, TM2 cell clusters, AQP1 was confined to TM2 and smooth muscle cell cluster, and CRYAB was localized to all cell clusters with more preference to schwann cell-like, TM2, smooth muscle cell, melanocyte, and myelinating schwann cell clusters. TM cells in uveal/ corneoscleral meshwork secrete antithrombotic molecules(14) such as tissue plasminogen activator (PLAT) which helps in maintaining clear flow of aqueous through TM and SC regions. The expression of PLAT was primarily localized to lymphatic-vascular like endothelium and schwann cell-like cell clusters, with some TM1 and TM2 expression. Podoplanin (PDPN, aka D2-40) has been widely used as marker to detect lymphatic vessels and studies have reported that TM stains positive for D2-40 in histological sections(63). Consistent with this finding, in our scRNAseq, PDPN was confined to TM1 and TM2 cell clusters, but not the endothelial/lymphatic Schlemm’s canal.

**Figure 9:**
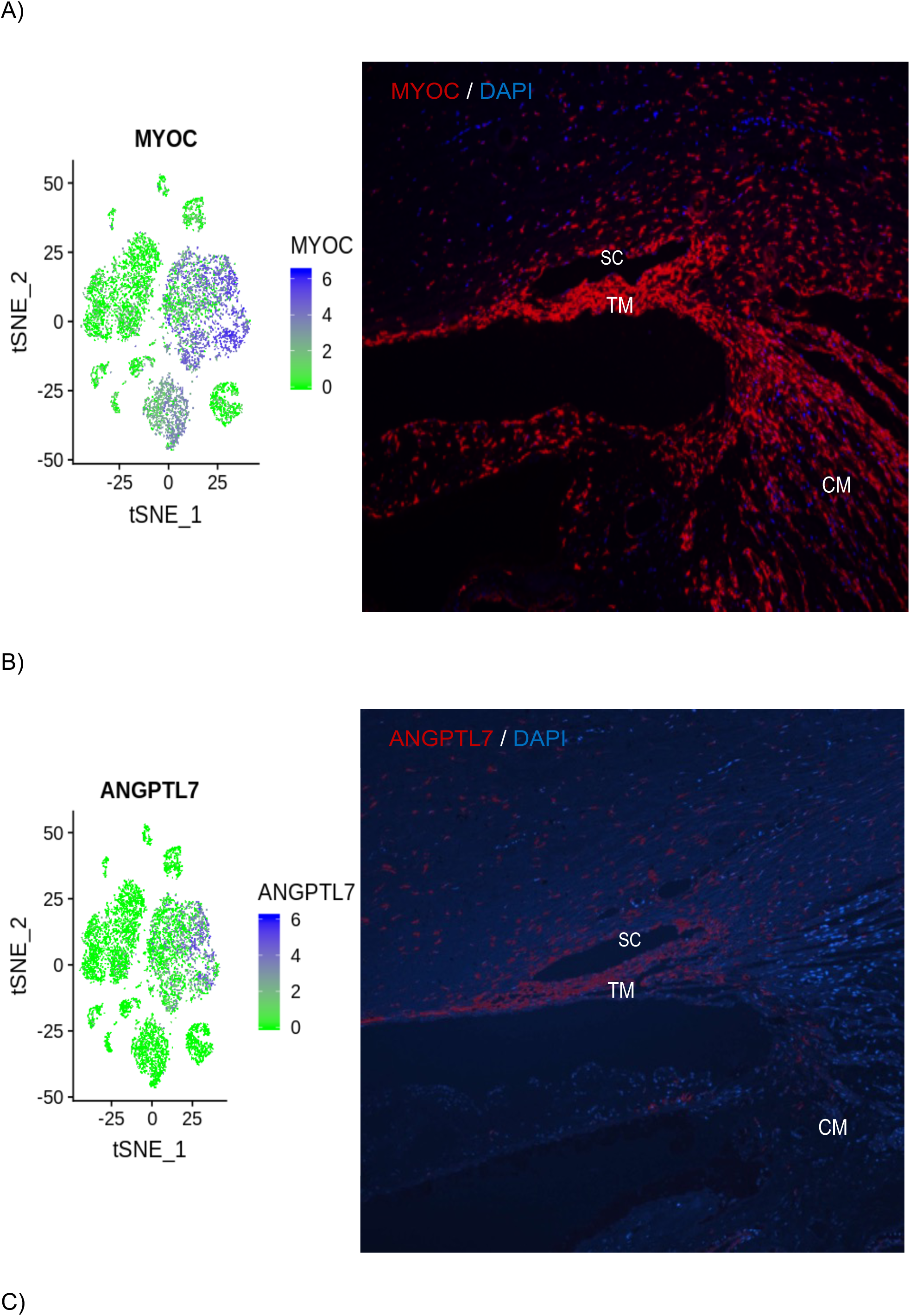
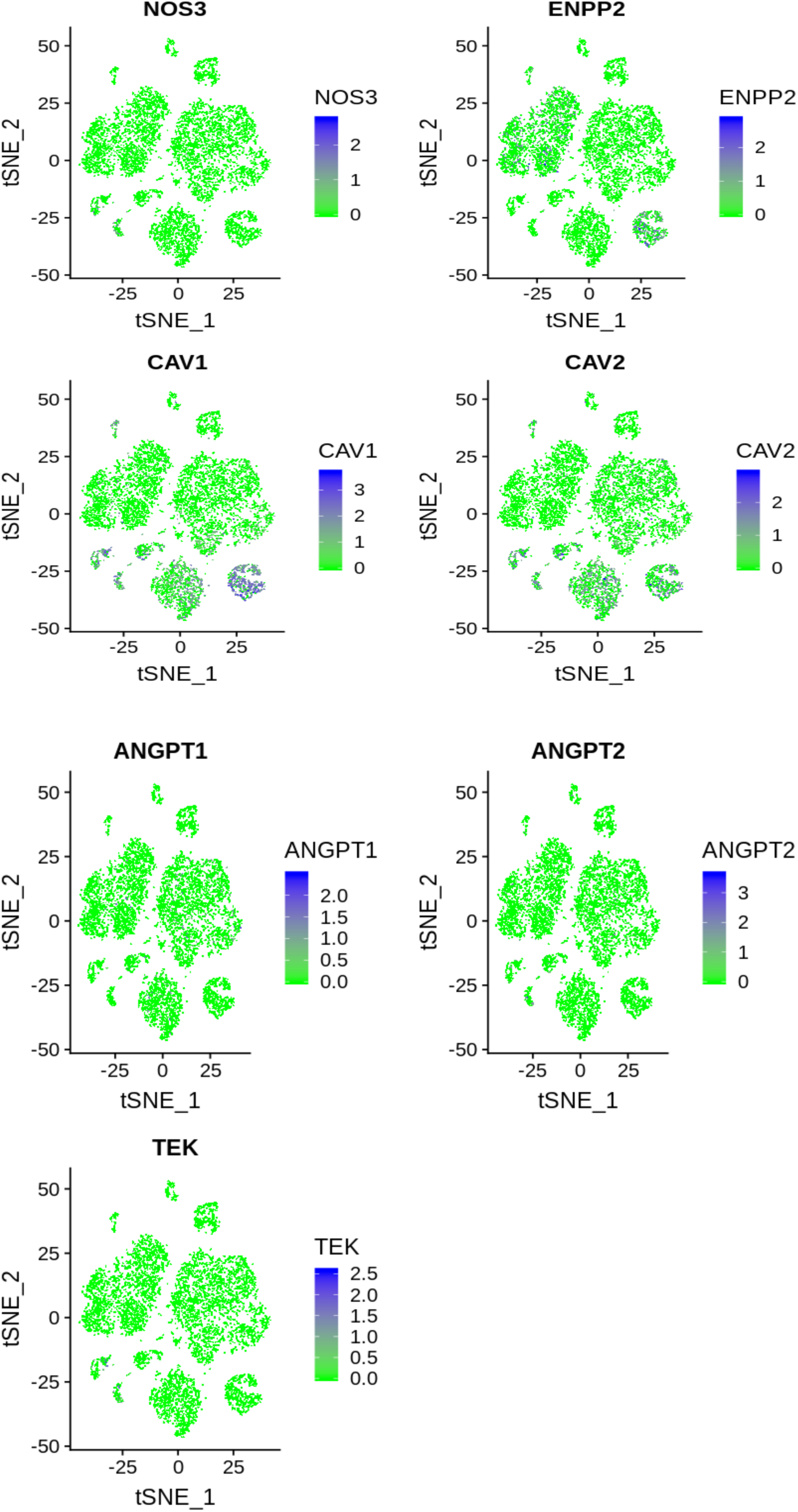
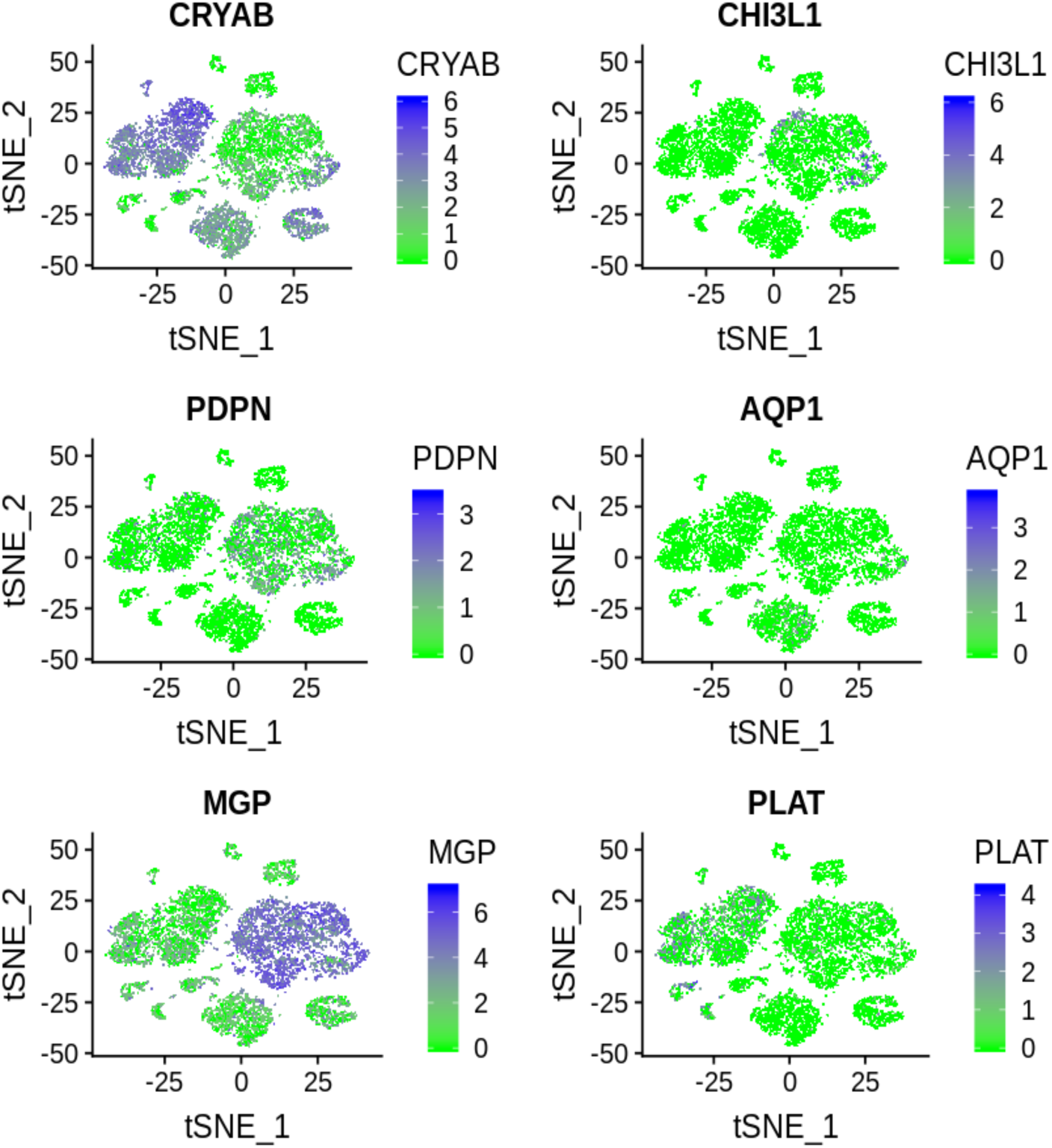
Expression of glaucoma-related genes in cell clusters identified in scRNAreq from human outflow cells and in human tissue sections by in situ hybridization. For panels A and B, the left side shows t-distributed stochastic neighbor embedding (tSNE) plot with relative expression of glaucoma gene (blue dots) in each cluster and right-side displays an in situ hybridization-stained human eye section showing mRNA signal as red dot fluorescence. mRNA probes specific for MYOC **(A)** localize strong expression throughout TM, with lower expression in the ciliary muscle and surrounding tissues. **(B)** ANGPTL7 was confined to JCT, SC region of the outflow tract in human eye sections. DAPI staining (blue) counterstains cell nuclei. Magnification: 20x. **(C)** T-distributed stochastic neighbor embedding (tSNE) plots with relative expression of glaucoma gene (blue dots) in each cluster. ENPP2, CAV1, CAV2, NOS3, ANGPT1, ANGPT2, TEK, CRYAB, CHI3L1, PDPN, AQP1, MGP, and PLAT gene expression in different cell clusters are displayed. TM- Trabecular meshwork; CM- ciliary muscle; SC- Schlemm’s canal. Scale in tSNE plot shows intensity of the average transcript count of particular marker within expressing cells and clusters.

**Table 3:** Glaucoma specific gene expression (average UMI) in different cell clusters.

| Gene | Lymphatic-like endothelium | Vascular endothelium | Pericyte | Smooth muscle cell | TM2, myofibroblast-like | TM1, fibroblast-like | Myelinating Schwann cell | Schwann cell-like | Melanocyte | Epithelium | T/NK cell | Macrophage |
| --- | --- | --- | --- | --- | --- | --- | --- | --- | --- | --- | --- | --- |
| NOS3 | 0.42 | 0.60 | 0.05 | 0.00 | 0.01 | 0.01 | 0.00 | 0.00 | 0.01 | 0.00 | 0.04 | 0.00 |
| ENPP2 | 0.00 | 0.10 | 0.03 | 0.03 | 0.02 | 0.07 | 0.53 | 0.69 | 1.60 | 0.00 | 0.02 | 0.05 |
| CAV1 | 5.17 | 2.92 | 2.32 | 1.76 | 0.59 | 0.11 | 2.75 | 0.14 | 7.35 | 0.62 | 0.30 | 0.09 |
| CAV2 | 2.14 | 2.09 | 1.62 | 1.40 | 0.41 | 0.27 | 1.31 | 0.09 | 1.79 | 0.86 | 0.08 | 0.04 |
| ANGPT1 | 0.01 | 0.02 | 0.16 | 0.09 | 0.12 | 0.11 | 0.01 | 0.02 | 0.00 | 0.00 | 0.00 | 0.00 |
| ANGPT2 | 0.13 | 0.59 | 0.33 | 0.02 | 0.05 | 0.07 | 0.08 | 0.06 | 0.04 | 0.03 | 0.09 | 0.01 |
| TEK | 0.84 | 0.62 | 0.04 | 0.00 | 0.01 | 0.01 | 0.01 | 0.00 | 0.00 | 0.00 | 0.00 | 0.00 |
| CRYAB | 1.77 | 1.35 | 8.20 | 15.66 | 14.03 | 3.52 | 70.56 | 50.57 | 33.77 | 4.24 | 1.32 | 1.59 |
| CHI3L1 | 0.02 | 0.12 | 1.42 | 0.08 | 10.48 | 4.34 | 0.15 | 0.16 | 0.05 | 0.11 | 0.27 | 0.17 |
| PDPN | 0.12 | 0.06 | 0.10 | 0.12 | 2.67 | 2.02 | 0.24 | 0.68 | 0.04 | 0.04 | 0.15 | 0.07 |
| AQP1 | 0.17 | 0.18 | 0.16 | 0.75 | 0.37 | 0.11 | 0.05 | 0.05 | 0.01 | 0.00 | 0.00 | 0.03 |
| MGP | 10.71 | 8.93 | 17.45 | 8.81 | 157.33 | 144.92 | 9.93 | 8.50 | 12.40 | 6.50 | 6.28 | 5.46 |
| PLAT | 12.78 | 2.68 | 0.28 | 0.10 | 0.34 | 0.49 | 1.34 | 2.80 | 0.31 | 0.77 | 0.08 | 0.02 |

### Immunohistochemical localization of candidate gene products in human outflow tissues

The protein products of select candidate genes identified by single cell RNAseq (table S1) were screened in sections of human outflow tissues. We focused on the four main cell types in the conventional outflow pathway, TM1, TM2, SC and macrophages. A positive controls for TM1 and TM2 was the glaucoma-related gene product myocilin, which labeled strongly in all regions of the TM, and have weaker labeling in surrounding tissues (Figure 10A). The vascular endothelial marker, PECAM1 (CD31) was used to selectively label SC as a positive control (Figure 10B). We tested a number of commercially-available antibodies at different concentrations against gene candidates, using a two different tissue preparation techniques on different donor eyes sections (Table S1, S2). We were unable to obtain specific labeling with a number of these antibodies, however we reliably observed labeling with the TM-1 candidate gene product, RSPO2 (Figure 10C). The labeling pattern was similar to that found with RNAscope (Figure 4), labeling both TM1 and TM2 cells, with preference to JCT (TM1 cells). Antibodies to its homolog, RSPO4 demonstrated a lot of non-specific labeling of TM and surrounding tissues (not shown). We also tried a number of different antibodies against SC candidates including endothelial nitric oxide synthase, VEGF receptor-3, trefoil factor-3 and TMEM88, but labeling was found to be non-specific or negative for SC and neighboring endothelial lined vessels. Lastly, we were interested in looking at candidate markers for myelin-containing neurons and resident macrophages and natural killer (NK) cells in the TM. For neurons, we labeled human outflow tissues with two different antibodies raised against myelin-PLP, without specific staining (not shown). For NK cells and macrophages, we labeled human outflow tissues antibodies raised against DAP12 and LYVE-1, respectively. Interestingly, the LYVE-1 staining was positive against some cells in the TM, with more prominent staining of the SC lumen and distal vessels (Figure 10D).

**Figure 10:**
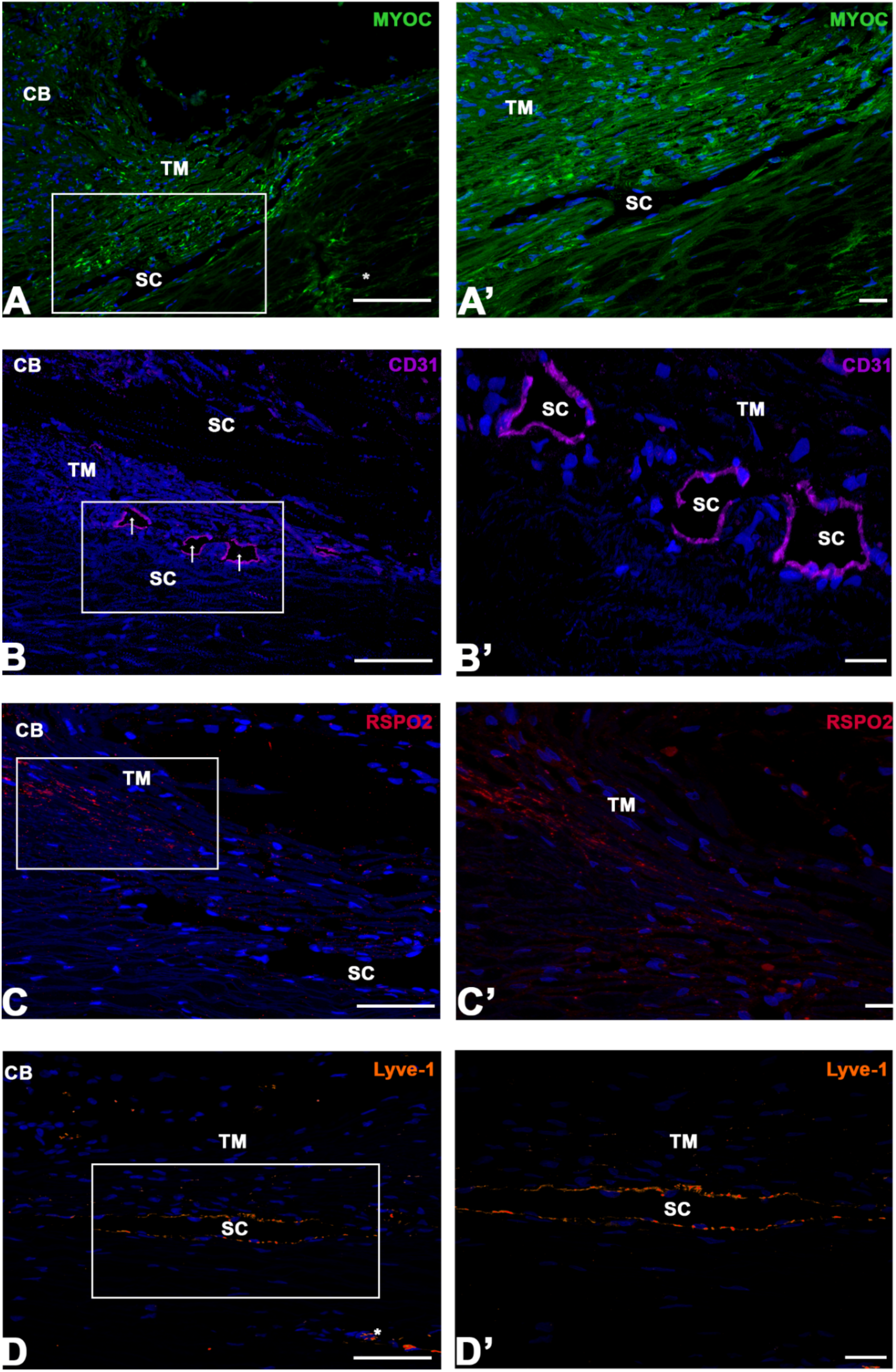
Immunolocalization of protein products of gene cluster markers in conventional outflow tissues from human eye sections. Left panels show low magnification view of conventional outflow structures in cross-section. Areas of interest are indicated by white box, which corresponds to higher magnification image in right panels. Panels A show strong trabecular meshwork (TM) labeling of the glaucoma gene, myocilin (MYOC). Panels B show staining for the vascular endothelial cluster marker, CD31 (PECAM1), which is limited to Schlemm’s canal. Panels C show the TM-2 gene cluster candidate, R-spondin-2 (RSPO2), displaying preferential labeling in the JCT region of the conventional tract. Panels D show the macrophage cluster marker, Lymphatic vessel endothelial hyaluronan receptor 1 (Lyve-1), showing predominant Schlemm’s canal and scleral vessel labeling. Asterisk indicates scleral vessel. CB: ciliary body, magnification bars in left panels = 100μm, and in right panels 20μm.

## DISCUSSION

The conventional outflow pathway is a complex tissue involved in maintaining IOP homeostasis. Cells that reside here have complementary roles to ensure faithful regulation of outflow resistance, and thus IOP. We used scRNAseq to generate an atlas of human conventional outflow cells to more accurately define their cellular roles based upon differential gene expression and tissue localization. The data set was robust, being generated from 8 individual samples from 4 human donor eyes. These expression data more accurately and convincingly identify the various individual cell types, cell-type specific gene markers, and disease-related gene expression across the different cell types in the conventional outflow pathway. Major outcomes of the present study were the identification of 12 different cell types in, or adjacent to conventional outflow tissues. Importantly, we clearly found that there are two distinct “molecular types” of TM cells with related expression patterns. We also showed in human eyes the hybrid blood vasculature/lymphatic character of SC, supporting mouse lineage tracing data. Finally, we demonstrate the utility of our data set by localizing select glaucoma-related genes to specific cell populations in the outflow tract. These findings have only tapped the surface of this valuable and reliable dataset that can be used by conventional outflow researchers for years to come.

One of the major cell types identified was the Schwann cell like cell cluster. Despite the abundance of this cell type in our single cell samples, it was found to not be a major complement of the TM proper. Our in-situ hybridization results mapped this cluster primarily to scleral spur and ciliary muscle region in human eye sections. These data are consistent with other studies showing the presence of myelinated nerves that innervate scleral spur cells, or mechanosensory nerve endings in the scleral spur and ciliary muscle region(8, 64–67). Electron micrographs of nerve terminals in the scleral spur show that the surface of such terminals is ensheathed by flat processes of Schwann cells(8, 67). The elastic fibers and nerve terminals of scleral spur are directly continuous with elastic fibers of TM and extend with ECM of TM(8, 65). Along this path, we found a few cells expressing these markers that extended into the TM (Figure 2), consistent with TEM data of human eyes(65).

Our results suggest that TM2, TM1, lymphatic-vascular endothelial cell and macrophage cell clusters represents the primary components of the conventional outflow pathway corresponding to TM cells on beams/cribriform plates, JCT cells, SC endothelium and macrophages throughout. While the candidate genes RSPO2 and RSPO4 were primarily found in the TM2-myofibroblast cell cluster, in situ hybridization demonstrated a broader distribution across the TM, with RSPO4 being more restricted to TM than RSPO2. Immunolabeling of RSPO2 protein confirmed in situ hybridization results, showing some preferential expression in TM cells. RSPO2 and −4 are secreted proteins that enhance wnt signaling, a known regulatory pathway of outflow resistance and IOP (68, 69). While also being expressed at low levels in neighboring tissues, the TM1-fibroblast cell cluster candidate genes DCN, PDGFRA preferentially localized to JCT region of outflow tract, a region critical for generating outflow resistance(7). TM cells in JCT region display fibroblast and smooth-muscle like phenotype and they modulate ECM turnover and repair(7). DCN encodes, decorin which is a proteoglycan that regulate ECM deposition and is an antagonist of TGF*β* and CTGF pathways. Diminished decorin levels have been observed in aqueous humor of glaucoma patients(70) and decorin treatment lowers IOP and RGC loss in animals models(71). Finally, the lymphatic-vascular endothelial cell cluster expressed markers for both vascular and lymphatic endothelial like phenotypes representing SC region which is unique vessel, having properties of both cell types(44). These primary data in human eyes support findings of mouse lineage tracing studies(44). The select group of glaucoma related genes we tested were found distributed to different outflow cell types in the conventional outflow pathway. For example, MYOC, ANGPTL7, PDPN, CHI3L1, and ANGPT1 were highly expressed in TM1, TM2 cell clusters, while CAV1, CAV2, NOS3, Tie2 (TEK), PLAT, and ANGPT2 were highly expressed in lymphatic-vascular endothelial cell clusters.

Unconventional outflow pathway structures which includes ciliary muscle and scleral spur cells were represented in our atlas by smooth muscle cell and scleral spur/Schwann cell like cell clusters. One of the candidate genes, TAGLN mapped to TM2 and smooth muscle clusters, appearing expressed at higher levels in ciliary muscle compared to TM by in situ hybridization. TAGLN encodes transgelin, which is an actin-binding protein that is ubiquitously expressed in vascular and visceral smooth muscle cells(72). These data emphasize the important contractile connection between the ciliary muscle and TM. Unexpectedly, autotaxin (gene name: ENPP2) which been shown to be elevated in aqueous humor of glaucoma patients(50, 51) was expressed in scleral spur/ Schwann cell like cluster and melanocyte cell clusters, but not TM1 or TM2. This is in contrast to previous reports finding autotaxin protein in human TM specimens(73). This highlights a limitation of current single cell technology, which is not sensitive enough to detect all the mRNA molecules in the cell(74). Very often mRNA of genes expressed at low level is not detected in every cell of the same cell type. This phenomena is called dropout, which is important to keep in mind when interpreting the biology of the scRNASeq data.

One of the important functions of TM cells is to clear debris delivered via aqueous humor flow as it moves across ocular structures in the posterior and anterior chambers. TM cells in the uveal/ corneoscleral meshwork have scavenger receptors and work as filter to clear cellular debris by phagocytosis before it reaches the resistance-generating JCT region (15–17). It appears that TM cells do not work alone but coordinate activities with resident macrophages as has been observed in smooth-muscle containing blood vessels(75). Consistent with this idea, our results map macrophage candidate genes (LYVE1, C1QB, TYROBP) abundantly throughout the conventional outflow tract, from inner TM to outer wall of SC and distal vessels, suggesting a homeostatic function in outflow regulation.

In all of our conventional outflow samples we consistently identified cells that mapped to clusters including T/NK cell, epithelium, melanocyte and pericyte (Figure S5). We speculate that they may be contaminants introduced during the dissection, or from neighboring tissues. Interestingly, while T/NK, epithelium and pericyte were represented by few cells, a large number of cells appeared in the melanocyte cluster.

In conclusion, using sc-RNAseq we have identified 12 distinct cell types in and around the human conventional outflow pathway, emphasizing the diversity of cells that participate in the regulation of outflow function, and thus IOP control. In addition to documenting the unique lymphatic/blood vascular expression profile of SC, our findings also settle the long-standing controversy as to the number of TM cell “types” in the conventional outflow pathway, finding two. These data provide essential new information for identity confirmation of TM and SC cells in culture, for designing cell specific promoters and for testing druggable targets for novel glaucoma therapies.

## MATERIALS AND METHODS

### Human eye tissue procurement and dissection

Human donor eyes (4 normal donors) were obtained from the Lions Eye Institute for Transplant and Research (Tampa, FL) and Miracles in Sight (Winston-Salem, NC) in moist chamber on ice (Table 1). The whole globes were soaked in betadine solution for 2-3 minutes and then rinsed with PBS twice. Using scalpel blade incision was 1mm posterior of limbus and eye globe was cut open with curved scissors along the circumference keeping 1mm distance from limbus, sometimes requiring cutting through vitreous. Using forceps, anterior and posterior eye globes were separated and from anterior part of eye, lens, iris, ciliary body were gently pulled off. Iris/pigment, ciliary muscle, and other tissue debris were gently scrapped off using edge of a clean blade and brief rinse of tissue in PBS. For dissecting trabecular meshwork (TM), blunt dissection approach was used(26). This was accomplished by lifting TM with forceps and teasing away continuous strand of TM tissue between the Schwalbe’s line and the scleral spur. The TM tissue was placed in digestion buffer containing 5mg collagenase A (Worthington Biochemical Corporation, Lakewood Township, NJ) dissolved in human albumin (catalog#A9080, Sigma-Aldrich). H&E staining of TM tissue before and after dissection is shown in supplementary figure S1A and S1B.

**Table 1:**
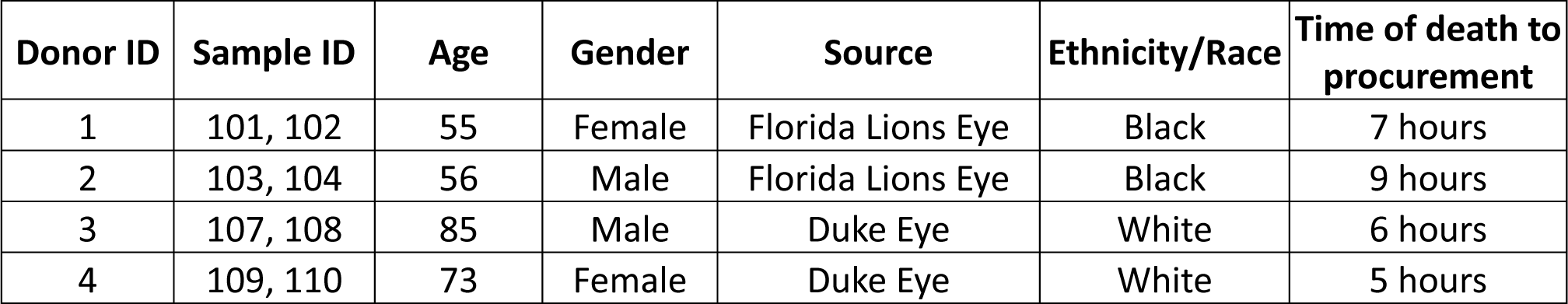
Sample description (sources, donor details)

| Donor ID | Sample ID | Age | Gender | Source | Ethnicity/Race | Time of death to procurement |
| --- | --- | --- | --- | --- | --- | --- |
| 1 | 101, 102 | 55 | Female | Florida Lions Eye | Black | 7 hours |
| 2 | 103, 104 | 56 | Male | Florida Lions Eye | Black | 9 hours |
| 3 | 107, 108 | 85 | Male | Duke Eye | White | 6 hours |
| 4 | 109, 110 | 73 | Female | Duke Eye | White | 5 hours |

### Sample preparation (cell dissociation and viability test)

The TM tissue in digestion buffer were incubated for 2-3 hours at 37°C and 5% CO_2_, shaking every 20 minutes. At the end of incubation, TM single cell suspension and tissue debris were mixed with media containing Dulbecco’s modified Eagle’s medium (DMEM) (Invitrogen-Gibco Life Technologies, Grand Island, NY, USA) supplemented with 10% fetal bovine serum (FBS; Atlas Biologicals, Fort Collins, CO, USA), penicillin (100 units/mL), streptomycin (0.1 mg/ mL), and L-glutamine (0.292 mg/mL) (Thermo Fisher Scientific, Rockford, IL, USA). The single cell suspension solution was filtered through 70-micron filter retaining tissue debris on filter. The single cell solution was centrifuged at 1000g for 10 minutes. The single cell pellet was dissolved in PBS with 0.04% BSA. Cell viability was determined by automated cell analyzer NucleoCounter® NC-250™.

### Single cell RNA sequencing, read mapping

Single cells suspended in PBS with 0.04% BSA were loaded on a Chromium Single Cell Instrument (10X Genomics). RNAseq libraries were prepared using Chromium Single Cell 3’ Library, Gel Beads & Multiplex Kit (10X Genomics). To minimize the presence of doublets in our population, approximately 6000 cells were loaded per lane and 2000-3000 cells per lane were recovered. Paired-end sequencing was performed on Illumina NextSeq500 (Read 1 26-bp for unique molecular identifier (UMI) and cell barcode, 8-bp i7 sample index, 0-bp i5, and Read 2 55-bp transcript read). Cell Ranger Single-Cell Software Suite (10X Genomics, v2.0.0) was used to perform sample de-multiplexing, alignment, filtering, and UMI counting. Human b37.3/ Mouse mm10 Genome assembly and UCSC gene model were used for the alignment.

### Data Analysis

We mainly used Seurat 2.3 software package developed by Satija lab for the single cell data analysis. Seurat object was created using the DGE UMI data file for each of the 8 samples. Genes expressed in less than 5 cells in each sample were removed from analysis. Cells with number of genes detected in less than 500 or over 5000, or UMI ratio of mitochondria encoded genes vs. all genes over 0.20 were also removed. Data normalization and scaling for each cell were achieved by using Seurat global-scaling “LogNormalize” method, which normalizes the gene expression measurements by the total expression followed by multiplying a scaling factor of 10,000 and log-transformation. To avoid potential sample-to-sample variation caused by technical variation at various experiment steps, we employed Seurat data integration method. First, top 1000 variable genes of each of the 6 Seurat objects were identified using “FindVariableGenes”. The union of these variable genes that were detected in at least 5 cells in each of the 6 samples were used for Seurat “RunCCA” function of canonical correlation analysis. Subsequently, 13 dimensions were used to run AlignSubspace to combine the 6 Seurat objects into 1. Cells were then grouped into clusters by using Seurat functions “RunTSNE” and “FindClusters” of resolutions 0.8-1.0. Marker genes for each cluster were identified using Seurat function “FindAllMarkers”. Parameters were used such that these genes were expressed in at least 25% of the cells in the cluster, and on average 1.28-fold higher than the rest of cells with a negative binomial test p value of less than 0.01. The expression of cluster marker genes as well as canonical cell type-specific genes were used to define the cell type for each cluster. The specificity of the canonical cell type-specific genes or cell cluster-specific genes were further examined by “VlnPlot”, “FeaturePlot” or “DotPlot” function of Seurat package. The similarity or dissimilarity among the identified cell types was examined by hierarchical clustering using Euclidean distance and complete linkage algorithm in R (R Core Team 2017, https://www.r-project.org/).

### In situ hybridization using *RNAScope*

The expression pattern of TM single cell cluster specific gene expression in the human donor eye was determined by in situ hybridization using RNAScope® according to manufacturer’s specifications (Advanced Cell Diagnostics). Briefly, 10% NBF fixed and paraffin embedded human donor eye cups were cut into 5 μm sections and mounted on SUPERFROST® Plus glass slides. For RNAScope, slides were baked on slide warmer for 1 hour at 60°C and deparaffinized. Tissue sections then underwent 10 minutes of Pretreat 1-using hydrogen peroxide treatment (ACD, 320037) at room temperature, followed by 20 minutes of boiling at 90°C in Pretreat 2-target retrieval buffer treatment (ACD, 320043) in Oster Steamer (IHC World, LLC, Model 5709) and 30 minutes of Pretreat 3-using protease plus treatment (ACD, 320037) at 40°C in a HybEZ Oven (ACD, 310010). Tissue sections were then incubated with DNase I for 15 minutes at 40°C to reduce potential background from probes binding to genomic DNA. Tissue sections were then washed five times with water, hybridized with RNAScope probes for target genes for 2 hours at 40°C and the remainder of the manufacturer’s assay protocol was implemented (ACD, 322360) from Amplified 1 to Amplified 6. The slides were washed twice (two minutes each at room temperature) with RNAScope wash buffer (ACD, 310091). Signal was detected by incubation with fast red working solution (1:60 ratio of Red B to Red A) at room temperature for 10 minutes in the absence of light, followed by washing the slides in water several times, mounting the slides and viewing under bright-field microscope. In some experiments, fluorescent signals were visualized and captured using an open-field Nikon Eclipse Ti-E microscope.

### Immunohistochemistry of human anterior segments

Radial wedges cut from anterior segments of human donor eyes (Table S1) were paraffin-embedded, sagittaly sectioned (5um), deparaffinized following xylene and 100% ethanol incubations, and antigen-retrieved in boiling citric acid buffer for 15 min. Blocking buffer was prepared with 5% normal serum, 1% bovine serum albumin, 0.025% triton-x in tris-buffered saline. Sections were incubated in blocking buffer at room temperature (RT) for 1 hour. Antibodies directed against protein targets (Table S2) were incubated with tissue sections in blocking buffer overnight at 4°C. Tissue on slides were coverslipped and merged z-stacks (∼10µm) were imaged at different magnifications using Nikon Eclipse confocal microscope. Controls were imaged at identical confocal settings as experimental.

## ACKNOWLEDGEMENTS

We thank Joshua R. Sanes and Tavé van Zyl for sharing results of their parallel study prior to the submission.

## SUPPLEMENTARY FIGURES AND TABLES

**Figure S1:**
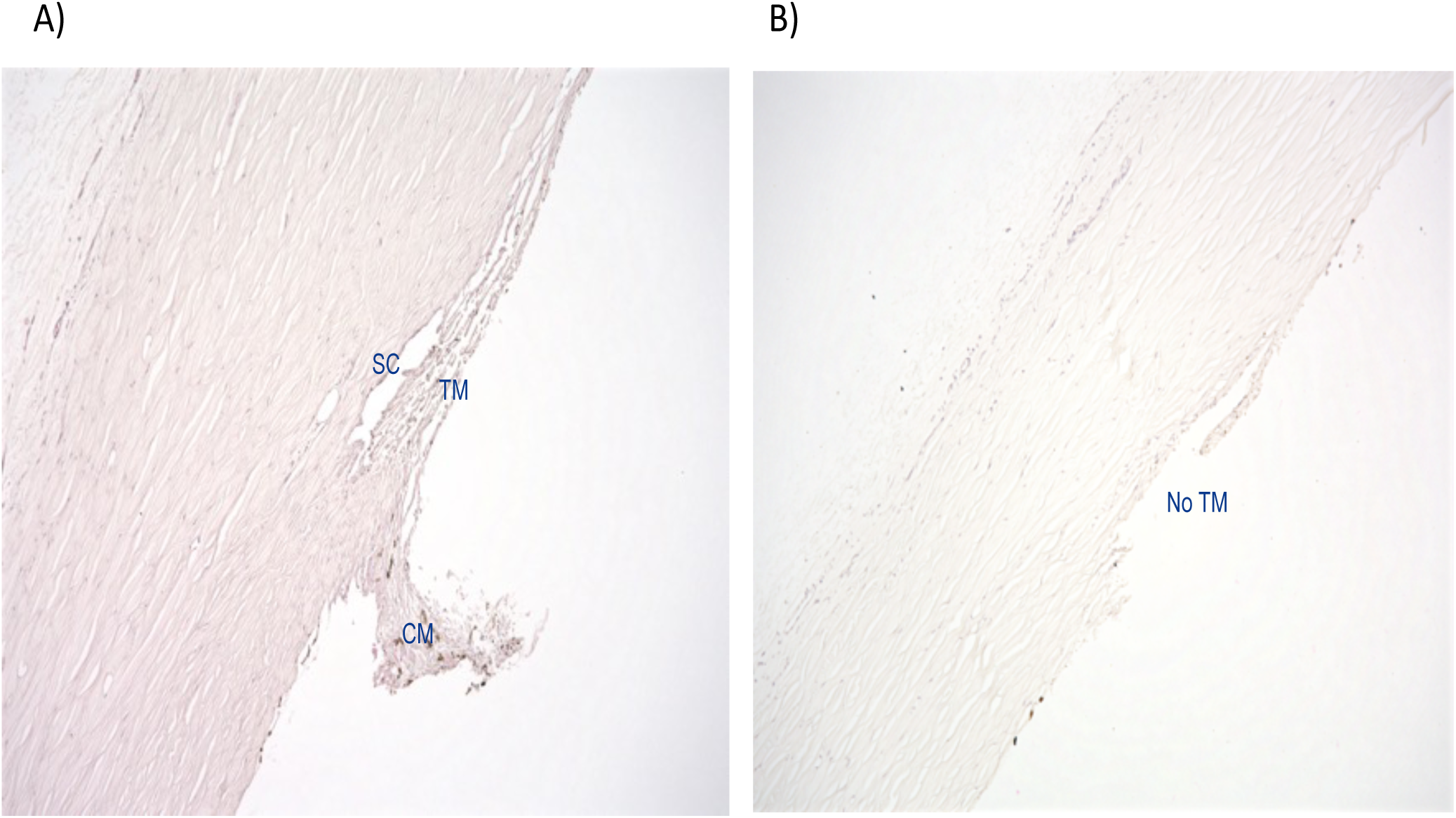
Blunt dissection of human outflow tissues. Hematoxylin and eosin stained human donor eye section showing intact TM before dissection **(A)** and after TM dissection **(B)**. Magnification: 20x. TM: Trabecular meshwork; SC- Schlemm’s canal; CM- Ciliary muscle

**Figure S2:**
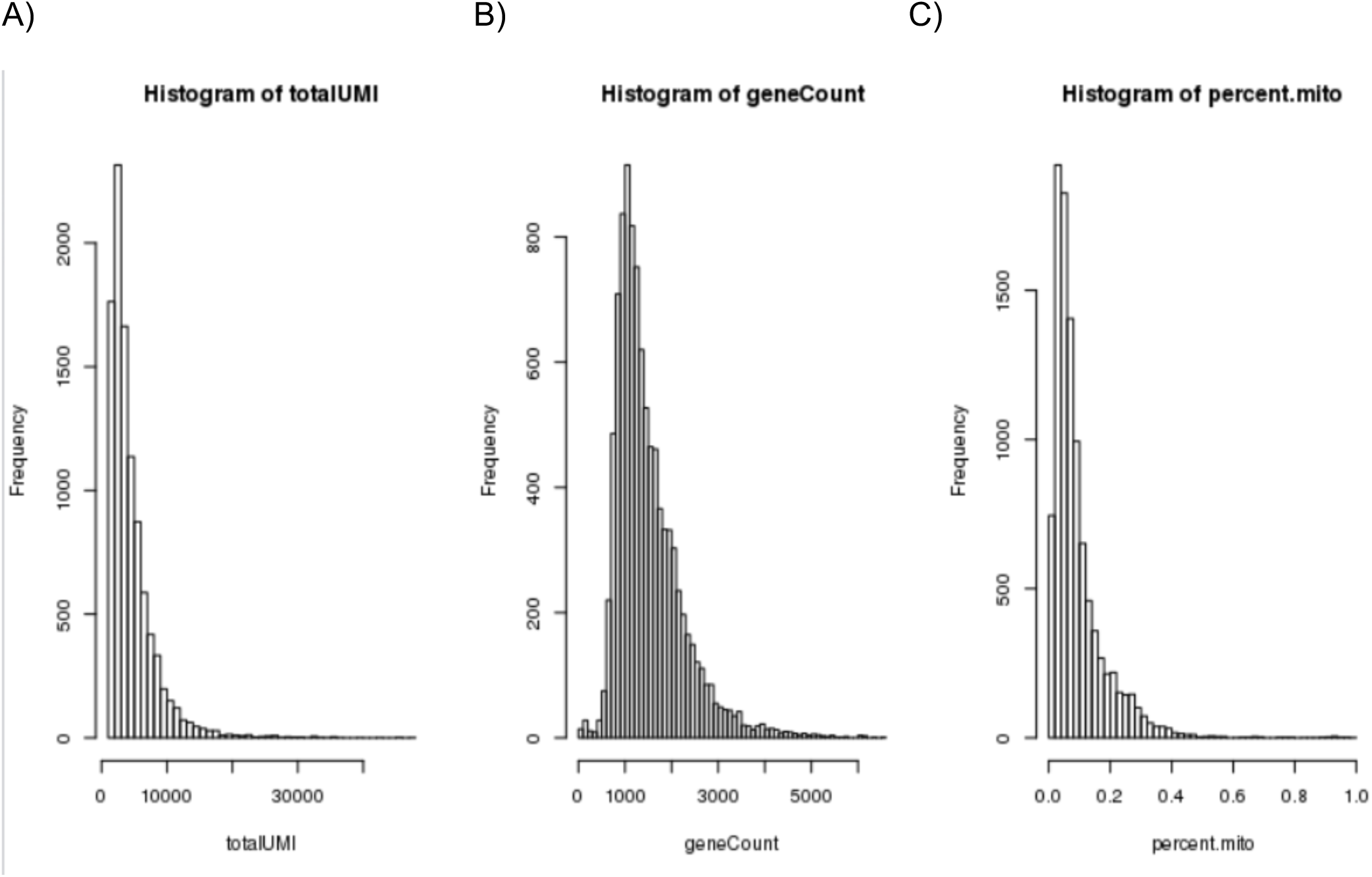
Three steps of QC. Histograms of total UMI (A), gene count (B) and percentage of mitochondria read counts (C)

**Figure S3:**
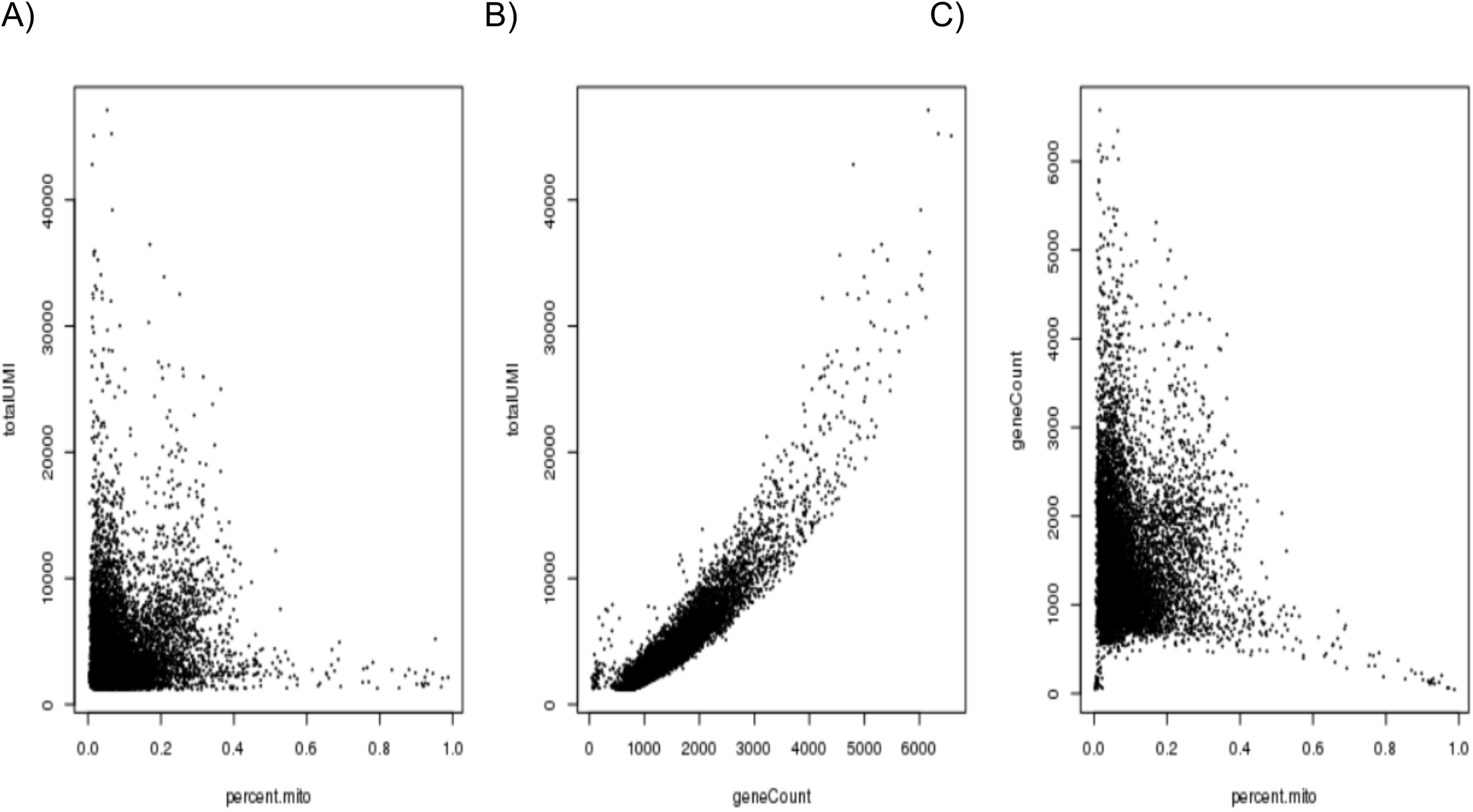
Scott plots of total UMI (A), gene count (B) and percentage of mitochondria read counts (C)

**Figure S4:**
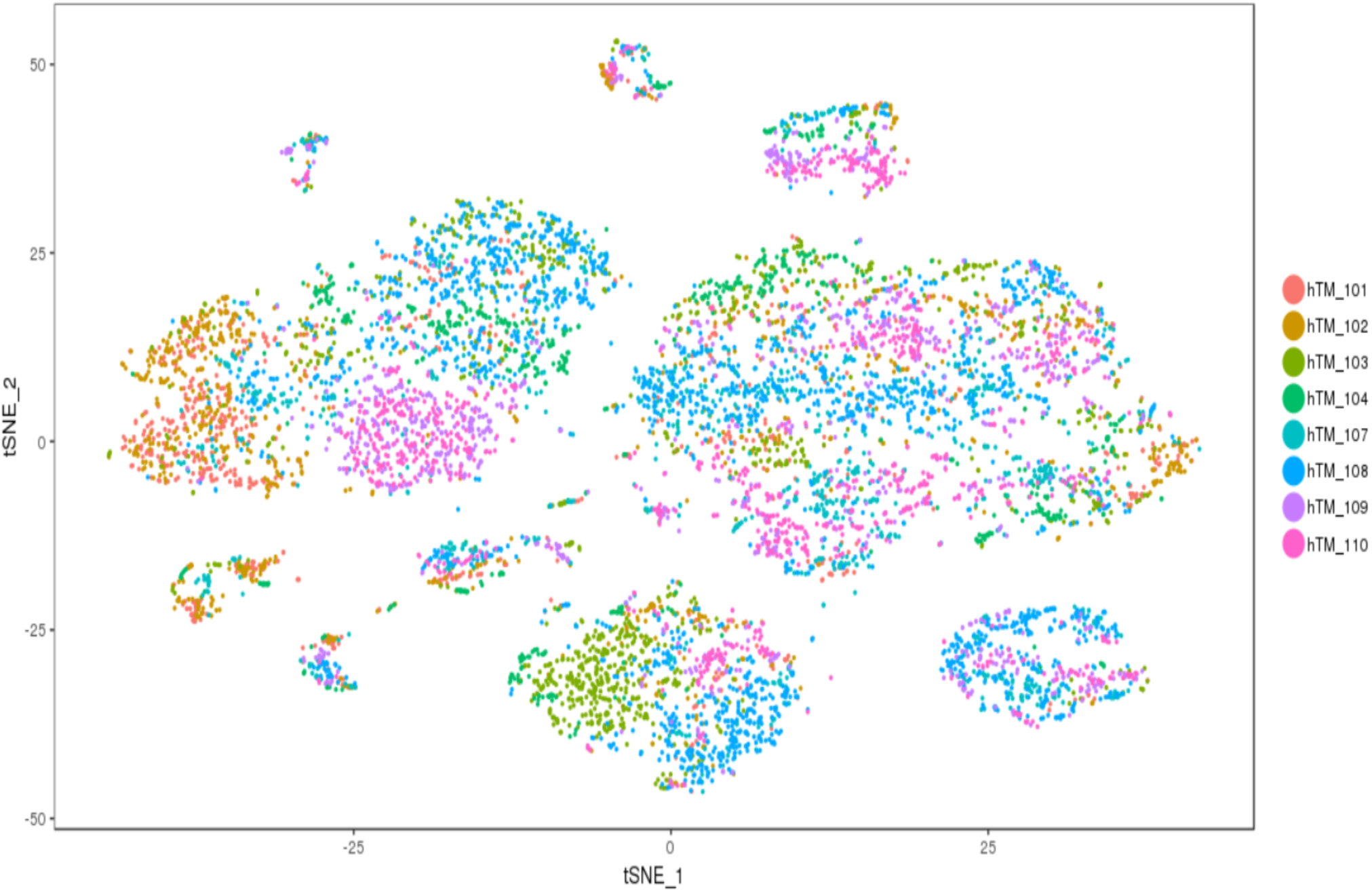
T-distributed stochastic neighbor embedding (tSNE) visualization of TM transcriptome heterogeneity of 8758 cells. Almost each of the human TM cell samples contributed uniformly to all clusters.

**Figure S5:**
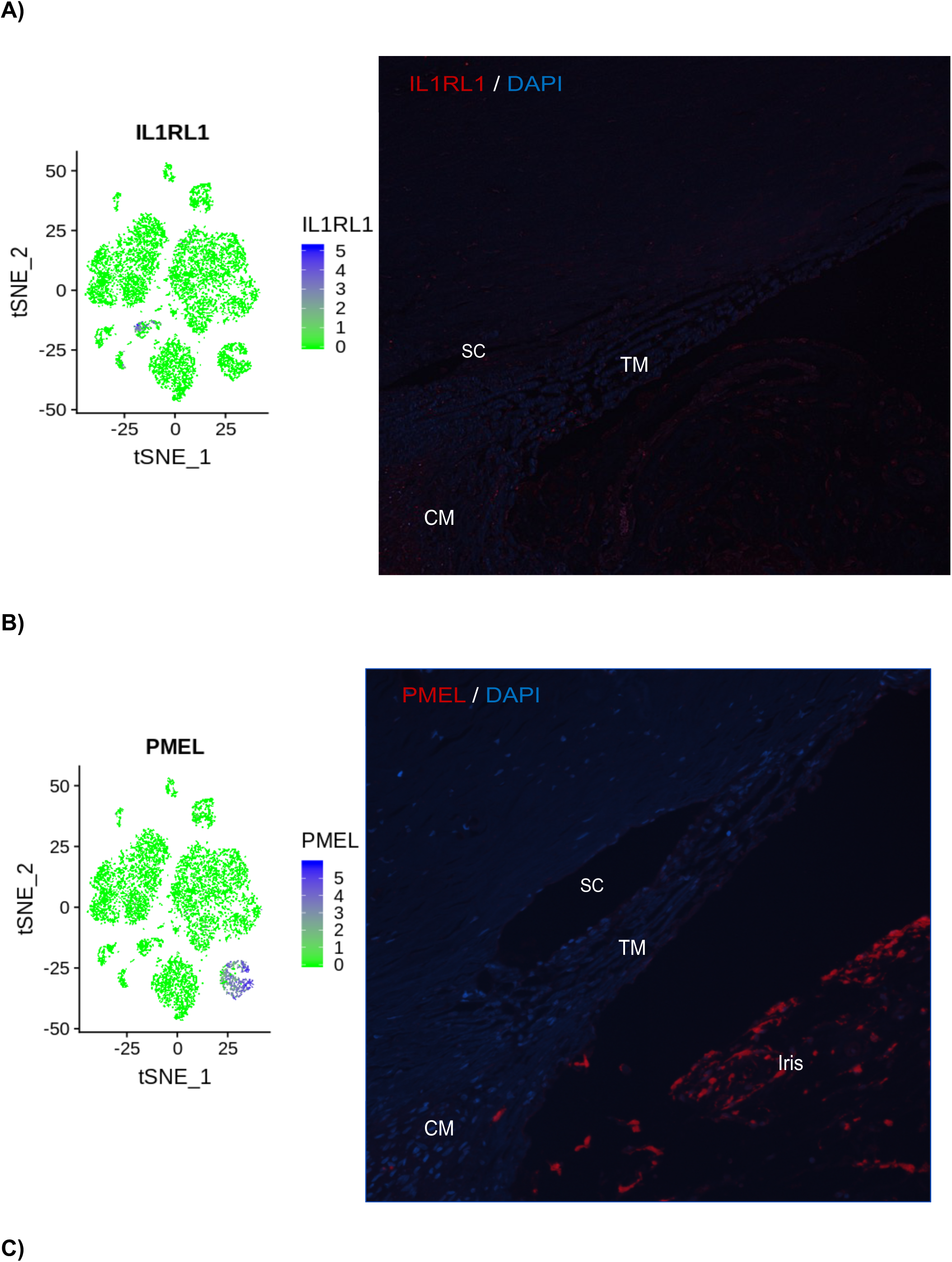
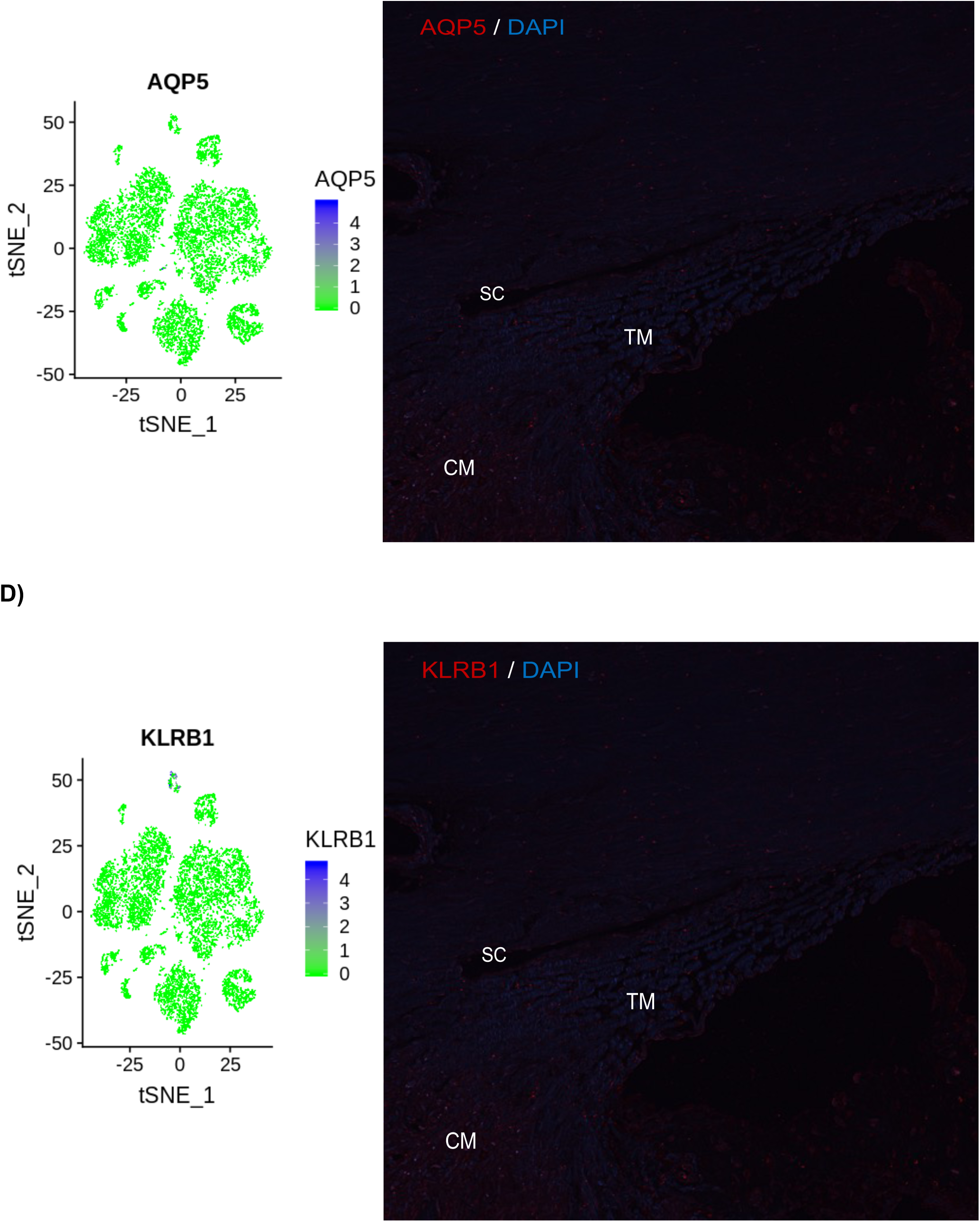
Mapping of pericyte, melanocyte, epithelium, and T/NK cell genes in human outflow tissues using in situ hybridization. For each panel, the left side is t-distributed stochastic neighbour embedding (tSNE) plot showing relative expression of gene of interest (blue dots) in each cluster and right-side is ISH stained human eye section showing mRNA signal as red dot fluorescence. mRNA probes to IL1RL1 **(A)**, PMEL **(B), (C)** AQP5, and **(D)** KLRB1 show that pericyte, melanocyte, epithelium, and T/NK cell are not present throughout the TM, ciliary muscle and around SC, respectively. DAPI staining (blue) counterstains cell nuclei. Magnification: 20x. TM- Trabecular meshwork; CM- ciliary muscle; SC- Schlemm’s canal. Scale in tSNE plot shows intensity of the average transcript count of particular marker within expressing cells and clusters.

**Supplementary table 1:** Candidate gene-products evaluated for expression in sagittal sections of human outflow tissues by immunohistochemistry.

| Target | Antibody | Predicted labeling | Identification numbers of donor eye tissues tested | Antibody dilutions tested | Frozen/Paraffin/Both | Staining Results |
| --- | --- | --- | --- | --- | --- | --- |
| CD31 | ab28364 | SC | 19000420-82 (OS and OD), 1900-0380-67 (OD and OS), 1900-1328-72 (OD and OS), 1900-1737 (OD and OS), CR1900-1971-38, CR1900-2037-50, 1900-0830 (OD only), 1800-2144 (OS), 1900-0731 (OS), 1900-0369 (OS) | 1:100, 1:50, 1:25 | B | Positive |
| MYOC | PMID: 9727403 | TM | 19000420-82 (OS and OD), 1900-0380-67 (OD and OS), 1900-1328-72 (OD and OS), 1900-1737 (OD and OS), CR1900-1971-38, CR1900-2037-50, 1900-0830 (OD only), 1900-0369 OS | 1:1000, 1:500, 1:250, 1200 | B | Positive |
| RSPO2 | EMD Millipore (MABS1709) | TM | 19000420-82 (OS and OD), 1900-0380-67 (OD and OS), 1900-1328-72 (OD and OS), 1900-1737 (OD and OS), CR1900-1971-38, CR1900-2037-50, 1900-0830 (OD only), 1800-2144 (OS), 1900-0731 (OS), 1900-0369 (OS) | 1:500, 1:250, 1:100, 1:50 | B | Positive |
| Lyve-1 | abcam ab14917 | Macrophage | 1900-0369 (OS) | 1:200, 1:100, 1:50 | P | Positive |
| RSPO4 | LS-C162784/143465 | TM | 19000420-82 (OS and OD), 1900-0380-67 (OD and OS), 1900-1328-72 (OD and OS), 1900-1737 (OD and OS), CR1900-1971-38, CR1900-2037-50, 1900-0830 (OD only), 1800-2144 (OS), 1900-0731 (OS) | 1:500, 1:250, 1:100, 1:50, 1:25, 1:15, 1:10 | B | Negative |
| Chromogranin B | ab1242 | TM | 1900-0830 (OD only), 1800-2144 (OS), 1900-0731 (OS), 1900-0369 (OS) | 1:1000 -1:100 | P | Negative |
| VEGFR3 | AF349 | SC | 1900-0830 (OD only), 1800-2144 (OS), 1900-0731 (OS), 1900-0369 (OS) | 1:500 -1:25 | P | Negative |
| eNOS | Novus NB300-500 | SC | 1900-0369 (OS) | 1:50-1:250 | P | Negative |
| eNOS | abcam5589 | SC | 19000420-82 (OS and OD), 1900-0380-67 (OD and OS), 1900-1328-72 (OD and OS), 1900-1737 (OD and OS), CR1900-1971-38, CR1900-2037-50, 1900-0830 (OD only), 1800-2144 (OS), 1900-0731 (OS) | 1:100 -1:15 | B | Negative |
| Trefoil Factor 3 | Origene AM20949PU-N | SC | 19000420-82 (OS and OD), 1900-0380-67 (OD and OS), 1900-1328-72 (OD and OS), 1900-1737 (OD and OS), CR1900-1971-38, CR1900-2037-50, | 1:500 -1:15 | B | Negative |
| TMEM88 | PA5-21164 | SC | 19000420-82 (OS and OD), 1900-0380-67 (OD and OS), 1900-1328-72 (OD and OS), 1900-1737 (OD and OS), CR1900-1971-38, CR1900-2037-50, 1900-0830 (OD only), 1800-2144 (OS), 1900-0731 (OS), 1900-0369 (OS) | 1:500 -1:25 | B | Negative |
| DAP12 | Origene AM20949PU-N | NK cells | 19000420-82 (OS and OD), 1900-0380-67 (OD and OS), 1900-1328-72 (OD and OS), 1900-1737 (OD and OS), CR1900-1971-38, CR1900-2037-50, 1900-0830 (OD only), 1800-2144 (OS), 1900-0731 (OS), 1900-0369 (OS) | 1:500 -1:25 | B | Negative |
| **DAP12** | BP1-85313 | NK cells | 1900-0830 (OD only), 1800-2144 (OS), 1900-0731 (OS) | 1:25-1:250- | P | Negative |
| **Myelin PLP** | ab28486 | Schwann | 1900-0830 (OD only), 1800-2144 (OS), 1900-0731 (OS), 1900-0369 (OS) | 1:500 -1:25 | P | Negative |
| **Myelin PLP** | ab183493 | Schwann | 19000420-82 (OS and OD), 1900-0380-67 (OD and OS), 1900-1328-72 (OD and OS), 1900-1737 (OD and OS), CR1900-1971-38, CR1900-2037-50, 1900-0830 (OD only), 1800-2144 (OS), 1900-0731 (OS), 1900-0369 (OS) | 1:500 -1:25 | B | Negative |

TM: trabecular meshwork; SC: Schlemm's canal; CD31: PECAM1; MYOC: myocilin; F: frozen sections; P: paraffin sections; B: both.

**Supplementary table 2:** Human donor eye tissue tested for expression of candidate gene products.

| Human Donor ID | Age | Race | Sex | Ocular history | Death to preservation |
| --- | --- | --- | --- | --- | --- |
| **1900-0420** | 82 YO | W | F | No ocular history | 6 hrs |
| **1900-0380** | 67 YO | W | F | No ocular history | 6 hrs |
| **1900-1328** | 72 YO | AA | F | No ocular history | 6.5 hrs |
| **1900-1737** | 57 YO | W | M | No ocular history | 6 hrs |
| **1900-1971\*** | 38 YO | W | M | No ocular history | 2.5 hrs |
| **1900-2037\*** | 50 YO | W | F | No ocular history | 8.5 hrs |
| **1800-2144** | 72 YO | W | F | Pseudophakic | 7 hrs |
| **1900-0731** | 58 YO | W | M | No ocular history | 6 hrs |
| **1900-0731** | 58 YO | W | M | No ocular history | 6.20 hrs |
| **1900-0369** | 84 YO | AA | M | Pseudophakic, dry eye | 5.3 hrs |

\*corneal rims, W: white, AA; African American; F: female; M: male

